# Glioblastoma remodeling of neural circuits in the human brain decreases survival

**DOI:** 10.1101/2021.02.18.431915

**Authors:** Saritha Krishna, Abrar Choudhury, Kyounghee Seo, Lijun Ni, Sofia Kakaizada, Anthony Lee, Alexander Aabedi, Caroline Cao, Rasika Sudharshan, Andrew Egladyous, Nyle Almeida, Humsa S. Venkatesh, Anne Findlay, Srikantan Nagarajan, David Raleigh, David Brang, Michelle Monje, Shawn L. Hervey-Jumper

## Abstract

Gliomas synaptically integrate into neural circuits. Prior work has demonstrated bidirectional interactions between neurons and glioma cells, with neuronal activity driving glioma growth and gliomas increasing neuronal excitability. In this study we wanted to know how glioma induced neuronal changes influence neural circuits underlying cognition and whether these interactions influence patient survival. We use intracranial brain recordings during lexical retrieval language tasks in awake humans in addition to site specific tumor tissue biopsies and cell biology experiments. We find that gliomas remodel functional neural circuitry such that task-relevant neural responses activate tumor-infiltrated cortex, beyond cortical excitation normally recruited in the healthy brain. Site-directed biopsies from functionally connected regions within the tumor are enriched for a glioblastoma subpopulation that exhibits a distinct synaptogenic and neuronotrophic phenotype. Tumor cells from functionally connected regions secrete the synaptogenic factor thrombospondin-1, which contributes to the differential neuron-glioma interactions observed in functionally connected tumor regions compared to tumor regions with less functional connectivity. The degree of functional connectivity between glioblastoma and the normal brain negatively impacts both patient survival and language task performance. These data demonstrate that high-grade gliomas functionally remodel neural circuits in the human brain, which both promotes tumor proliferation and impairs cognition.

## Main Text

Gliomas, including glioblastoma, exist within the context of complex neural circuitry. Neuronal action potentials promote glioma growth through both paracrine signaling (Neuroligin-3 and Brain Derived Neurotrophic Factor, BDNF) and AMPAR- mediated excitatory electrochemical synapses^1–4^. Likewise, glioblastomas influence neurons to induce neuronal hyperexcitability through non-synaptic glutamate secretion^5, 6^, secretion of synaptogenic factors^7^, and reduced inhibitory interneuron numbers within tumor-infiltrated cortical regions^8^. Beyond preclinical models, we previously demonstrated in awake, resting patients that glioblastoma-infiltrated cortex exhibits increased neuronal excitability^3^. It is therefore possible that glioma synaptic integration into neural circuits and glioma-induced neuronal changes may make tumor-infiltrated brain physiologically disorganized and dysfunctional during cognitive tasks. Alternatively, it is possible that glioma-neuron interactions may remodel functional neural circuits such that cognitive tasks stimulate the neuron-glioma network. If the latter were true, the functional connectivity of glioma may ultimately influence patient survival.

To address these hypotheses, we studied clinical, cognitive and electrophysiological neural responses in adult patients with newly diagnosed high-grade gliomas, together with cellular and molecular neuron-glioma interactions, in order to understand the influence of gliomas on neural networks (Fig. 1a-d, Extended Table 1).

**Fig. 1.**
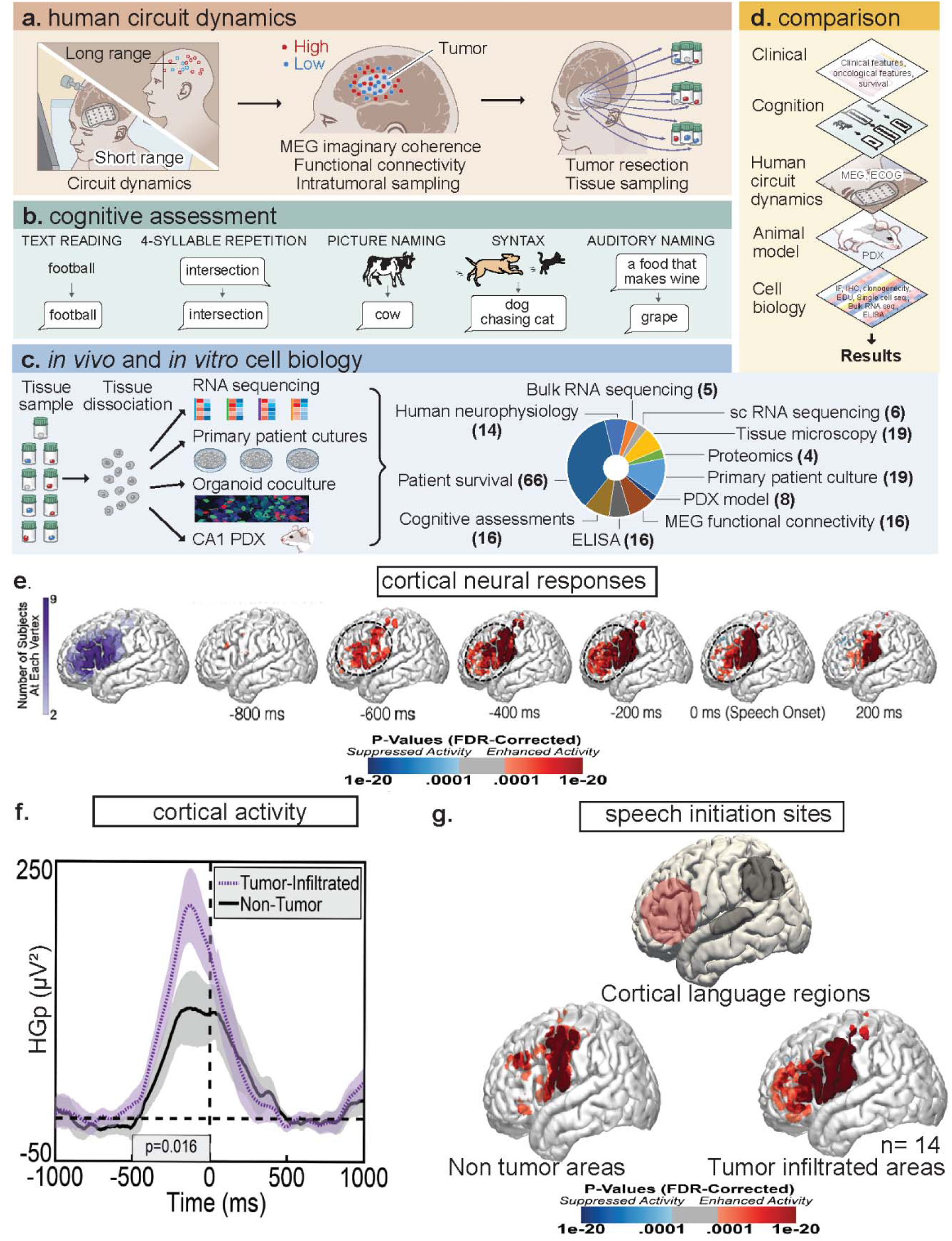
High-grade gliomas remodel long-range functional neural circuits. a, Schematic of study workflow. In human subjects with dominant hemisphere gliomas we applied electrocorticography during audiovisual speech initiation task to assess circuit dynamics. Focusing on isocitrate dehydrogenase wild type (IDH-WT) glioblastoma, we then applied a long-range measure of functional connectivity using magnetoencephalography (MEG) imaginary coherence. b, Extra- operative language assessments for correlation with biological assays. c, Long-range measure of functional connectivity to identify intratumoral regions of high and low functional connectivity for site specific biopsies which were used for *in vivo* and *in vitro* cell biology experiments. d, Multiple layered approach including clinical variables, cognition assessments, human and animal model network dynamics, and cell biology experiments serves as a platform for glioma influence on neural circuit dynamics. e, Electrocorticography electrodes placed within the dominant hemisphere speech initiation region of the posterior lateral frontal cortex (outlined area) of patients with high-grade gliomas (purple). Time series of human subjects picks up high gamma band power (HGp) within tumor infiltrated brain in 14 human participants between -600 milliseconds (ms) and speech onset (0 ms). f, Electrodes were compared between non-tumor and tumor infiltrated regions, FDR corrected HGp demonstrates task relevant hyper excitability (P= 0.016). g, These data illustrate task specific, speech initiation circuit remodeling within cortical language regions.

### Gliomas remodel functional neural circuits

High-grade gliomas interact with normal neuronal elements, resulting in both cellular and network level changes^9–11^. While high-grade gliomas influence neuronal excitability at rest, the effects of task-related activity on glioma-infiltrated neural circuit function and the impact of glioma-neuron interactions on neural circuit connectivity remain unknown. To examine cognitive task-related neuronal activity in the setting of high-grade glioma, we selected a cohort of 14 adult patients with cortically projecting glioma infiltration in the lateral prefrontal cortex (LPFC), classically referred to as Broca’s area (Extended Data Fig.1). In the operating room, tumor boundaries were localized on magnetic resonance imaging (MRI) and electrocorticography (ECoG) electrodes placed over the tumor-infiltrated cortical region and normal-appearing cortex. ECoG signals filtered between 70-110 Hz were used for analysis of the high-gamma band range power (HGp), which is strongly related to local neuronal population spikes^12, 13^ and is increased by cortical hyperexcitability^14^.

ECoG was recorded from the dominant hemisphere LPFC during auditory and visual picture naming tasks as an illustrative example of a well-defined cognitive neuronal circuit with defined physiology^15^. In humans, speech initiation occurs in the LPFC (Broca’s speech area, Brodmann area 44)^15^. While patients were fully awake and engaged in these language tasks, HGp was recorded from single electrodes overlying tumor-infiltrated and normal-appearing regions of brain (Extended Data Fig. 2a). These recordings provide simultaneous high spatial and temporal resolution while sampling the neuronal population activity during auditory and visual initiation of speech within the LPFC.

Group-level HGp from non-tumor electrodes demonstrates the expected neural time-course within LPFC, showing activation anterior to primary motor cortex between 600 milliseconds (ms) before speech onset (0 ms), and maximal activation in motor cortex at speech onset (Extended Data Fig. 2b), consistent with prior established models of speech initiation previously demonstrated in non-human primates and humans^16, 17^. We then performed the same time series focused only on electrode arrays recording from tumor-infiltrated cortex. Countering the theory that glioma-synaptic integration may result in physiologically disorganized neural responses, we found task-relevant neural activity within the entire region of tumor-infiltrated cortex. Strikingly, this includes speech initiation-induced recruitment of not only LPFC, Broca’s region, as expected, but also regions of tumor-infiltrated cortex not normally involved in speech initiation (Fig. 1e). Taken together, these findings suggest that in subjects with high-grade gliomas affecting the dominant hemisphere LPFC, naming tasks induce physiologically organized neuronal activity within tumor-infiltrated cortex, well beyond the cortical territory normally recruited during this language task.

Task-evoked neural responses from tumor-infiltrated regions may be less than in healthy tissues. Thus, we wanted to understand whether the magnitude of task-related neural activity within tumor-infiltrated regions of brain oscillates similar to non-tumor regions. We therefore pair-matched each cortical electrode array (Extended Data Fig. 2c, d) which confirmed increased HGp within this expanded region of cortex infiltrated by tumor (*P* = 0.016) (Fig. 1f). These data further demonstrate that glioma integration into cortical regions results in functional neural circuit remodeling and task-specific hyperexcitability (Fig. 1g).

### Synaptogenic glioma cells are functionally organized

Having established that gliomas functionally remodel neuronal circuits, we next wanted to understand whether functional integration heterogeneity exists within a specific molecularly defined high-grade glioma subtype. Given the heterogeneity of glioblastoma subpopulations^18–20^ and our previous finding that oligodendrocyte precursor cell-like subpopulations are enriched for synaptic gene expression^3^, functionally connected regions may vary within tumors and differences in functional connectivity between tumor regions may be due at least in part to a subpopulation of glioma cells with differential synaptic enrichment. With the goal of sampling functionally connected regions within tumor, we measured neuronal oscillations within glioma-infiltrated brain by magnetoencephalography (MEG) and imaginary coherence functional connectivity for subjects with newly diagnosed IDH-WT glioblastoma^21–23^. The functional connectivity of an individual voxel was derived by the mean imaginary coherence between the index voxel and the rest of the brain^24^. Functional connectivity brain regions were separated into upper tertile (high connectivity, HFC) and low tertile (low connectivity, LFC) which permitted site-directed tumor sampling.

To identify biological drivers and molecular targets of neural circuits between functionally connected, HFC and non-functionally connected, LFC tumor regions, we performed bulk and single-cell RNA-seq from tumor samples from each. Bulk RNA-seq transcriptomic analysis revealed upregulation in HFC tumor regions of a number of genes involved in neural circuit assembly, including axon pathfinding genes (*Netrin G1, NTNG1*), synapse-associated genes (e.g. *Synaptopodin, SYNPO*) and synaptogenic factors including a robust, 7-fold upregulation of *thrombospondin-1* (*TSP-1*). *TSP-1* was particularly intriguing in the context of the observed remodeling of neuronal network connectivity described above, because it is a known synaptogenic factor secreted by healthy astrocytes in the normal central nervous system^25^ (Fig. 2a, Extended Data Fig. 3a, b).

**Fig. 2.**
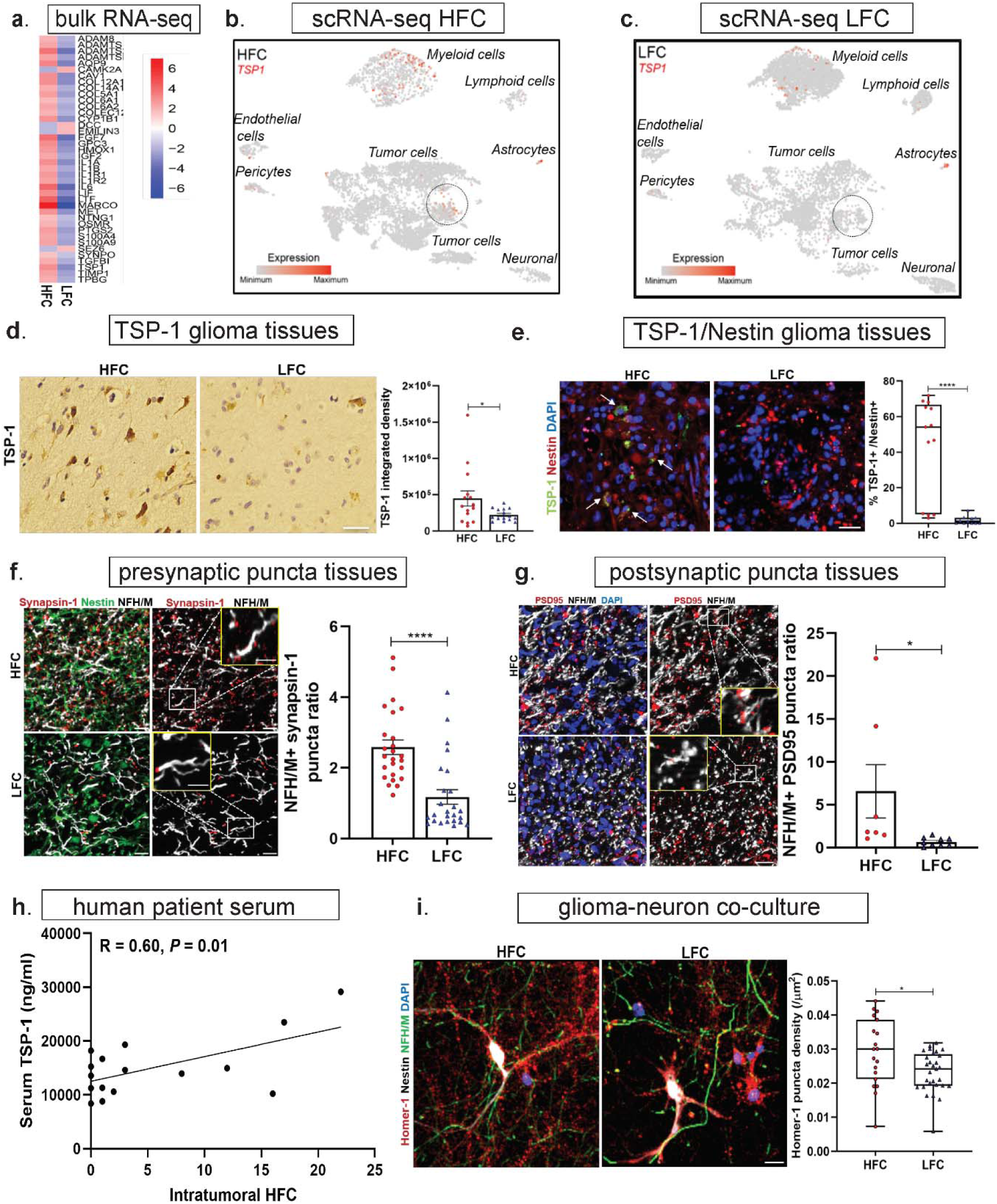
Tumor-infiltrated circuits exhibit areas of synaptic remodeling characterized by glioma cells expressing synaptogenic factors. a, Bulk RNA transcriptomic profile of HFC tissues showed a neurogenic signature including elevated (7-fold) expression of thrombospondin-1 (*TSP-1*) (n = 3-4 per group). b, c, Feature plots for *TSP-1* in HFC (b) (n = 6,666 cells, 3 subjects) and LFC (c) (n = 7,065 cells, 3 subjects) populations; within HFC samples with *TSP-1* expression primarily in tumor cells (denoted by circled area), and within LFC samples with *TSP-1* expression primarily from non-tumor astrocytes (suggesting that within low connectivity intratumoral regions normal astrocytes secrete *TSP-1* to generate connectivity mirroring normal physiology, while within HFC regions tumor cells express *TSP-1* to generate connectivity). d, TSP-1 (immunohistochemistry) expression in arbitrary units (AU) in HFC and LFC tissues (HFC vs. LFC: 447,063.42 AU± 103,473.52 vs. 221,341.77 ± 21,633.86 AU) (n = 5-6 sections, n = 3 per group). (P=0.04) Scale bar, 50 µm. e, TSP-1 expression from nestin-positive tumor cells in HFC and LFC tissues (HFC vs. LFC: 42.90 ± 7.74% vs. 1.74 ± 0.68%) (n = 11-13 sections, n = 3 per group). (*P* = 0.000073). Scale bar, 50 µm. f, g, Immunofluorescence images of synaptic puncta in HFC and LFC tissues. Ratio of presynaptic synapsin-1 (f) puncta count on NFH/M+ neurons to total DAPI count (calculated by dividing the total number of synapsin-1 puncta on neurofilament-positive neurons to the total number of cells stained with DAPI in 135 x 135-μm field areas) in HFC and LFC glioblastoma tissue samples (HFC vs. LFC: 2.58 ± 0.21 vs. 1.18 ± 0.20; n = 4 per group, *P* = 0.000014). Red, synapsin-1 (presynaptic puncta); green, nestin (HFC/LFC-GBM cells); white, neurofilament (neurons); blue, DAPI. Scale bar, 10 µm. Inset shows zoomed-in view of synapsin-1 puncta on neurons (positive for neurofilament). Scale bar, 3 µm. Ratio of post-synaptic PSD95 puncta count (g) on NFH/M+ neurons cells to total DAPI count (calculated by dividing the total number of PSD-95 puncta on neurofilament-positive neurons to the total number of cells stained with DAPI in 135 x 135-μ areas) in HFC and LFC glioblastoma samples (HFC vs. LFC: 6.56 ± 3.12 vs. 0.67 ± 0.16); n = 7-9 sections per group, n =3 per group) (*P* = 0.04). Red, PSD95 (postsynaptic puncta); white, neurofilament (neurons); blue, DAPI. Scale bar, 10 µm. Inset shows zoomed-in view of PSD95 puncta on neurons (positive for neurofilament). Scale bar, 3 µm. h, Linear regression statistics illustrates that circulating TSP-1 (measured by ELISA) correlates with number of intratumoral high functional connectivity voxels (n = 16) (P=0.01). i, Primary patient-derived HFC and LFC cells co-cultured with mouse hippocampal neurons. Immunofluorescence images for postsynaptic marker Homer-1 (red) in 2-week co-culture of glioma cells (white; nestin) and mouse hippocampal neurons (green; neurofilament). Quantification of postsynaptic Homer-1 puncta density (calculated by dividing the number of puncta measured with the area of the image field) in HFC and LFC glioma cells (HFC vs. LFC: 0.0291 ± 0.0023 µm^-2^ vs.0. 0232 ± 0.001 µm^-2^; n=4/group) (P=0.01). Scale bar, 50 μm. Data presented as mean ± s.e.m (d-g, i). *P* values determined by two-tailed Student’s t-test. \**P* < 0.05. \*\*\*\**P* < 0.0001.

To further determine whether any specific tumor cell population may be contributing to *TSP-1* expression, we performed single-cell sequencing of biopsy samples from HFC and LFC tumor regions (Extended Data Fig. 4a, b, Extended Data Tables 2, 3). Malignant tumor cells were inferred based on the expression programs and detection of tumor-specific genetic alterations including copy number variants (Extended Data Fig. 4c). We found that 2.4% of all tumor cells (HFC and LFC combined) expressed *TSP-1* and within this *TSP-1*-positive tumor cell population, HFC cells exhibit higher levels of *TSP-1* compared to LFC. Within LFC region samples, *TSP-1* expression originated primarily from a non-tumor astrocyte population (as determined by *S100* expression) (Extended Data Fig. 5). This data suggests that within low connectivity intratumoral regions, only non-tumor astrocytes express *TSP-1*, while within HFC regions, high-grade glioma cells express *TSP-1* in addition to non-tumor astrocytes which may promote increased connectivity (Fig. 2b, c, Extended Data Fig. 5). Elevated expression of *TSP-1* within HFC regions was confirmed by protein level analysis using HFC and LFC patient-derived glioblastoma biopsy tissues. Similar to our transcriptomic profiles, we found by immunohistochemistry increased TSP-1 expression within HFC tissues compared to LFC regions (Fig. 2d). Using immunofluorescence labelling we confirmed that malignant tumor cells in HFC regions are indeed TSP-1-expressing as determined by TSP-1/nestin co-localization within HFC tumor tissues (Fig. 2e). This finding that a subpopulation of malignant tumor cells in HFC regions produce TSP-1 suggests differential potential of tumor cells in the HFC regions to promote synaptogenesis and thereby connectivity. Glioblastomas are known for heterogenous cellular subpopulations which resemble both astrocyte and oligodendrocyte lineage^26, 27^, and previous studies have shown that astrocyte-like malignant cells can secrete synaptogenic factors that promote neuronal excitability^7, 28^. Our findings suggest that regions of tumor-infiltrated brain that exhibit increased functional connectivity include a synaptogenic subpopulation of malignant tumor cells. It is therefore possible that these synaptogenic glioma cells not only promote neuronal hyperexcitability but also potentially contribute to the functional circuit-level remodeling demonstrated above. These data are consistent with the cancer biology principal that cellular subpopulations assume distinct roles within the heterogenous cancer ecosystem which may be defined at least in part by functional connectivity measures.

Having established that high-grade gliomas can remodel functional circuits involved in language, that distinct intratumoral regions are enriched in functional connectivity and differentially contain a subpopulation of malignant cells expressing the synaptogenic factor TSP-1, we hypothesized that this subpopulation of HFC glioma cells may promote synaptogenesis and consequent remodeling of connectivity as observed in glioma-associated language networks above. To determine whether HFC-associated glioma cells promote structural synapse formation, similar to normal astrocytes^29–31^ and certain astrocyte-like glioblastoma cells^7, 28^, we first analyzed primary patient glioblastoma biopsies from HFC and LFC regions using immunohistochemistry and confocal microscopy. In these samples, neurofilament medium and heavy chain (NF) marks neuronal processes, synapsin-1 marks presynaptic puncta, and high-grade glioma cells are marked by nestin. We found increased presynaptic neuronal puncta (synapsin-1) within HFC regions with ∼ 2.58 synapsin-1 puncta per neuronal process compared with ∼1.18 synapsin-1 puncta per neuronal process within LFC region biopsies (P = 0.000014) (Fig. 2f). We similarly found increased postsynaptic puncta on neurons (PSD95-positive/neurofilament-positive) and glioma cells (PSD95-positive/nestin-positive) within HFC regions compared with LFC regions (Fig. 2g, Extended Data Fig. 6). We next examined whether TSP-1, a secreted synaptogenic protein^25, 29^, can be identified in patient serum and whether circulating TSP-1 is correlated with functional connectivity as measured by magnetoencephalography imaginary coherence. Circulating TSP-1 levels in patient serum exhibited a striking positive correlation with intratumoral functional connectivity (*P* = 0.01) (Fig. 2h), suggesting a role for TSP-1 in glioma-associated neural circuit remodeling and identifying a possible clinical correlate for functional connectivity in glioma patients.

We next created primary patient-derived glioma cultures from HFC and LFC tumor regions so that we could perform further mechanistic experiments and identify additional paracrine signaling molecules. Using these primary patient-derived cultures, we performed proteomics analysis of secreted factors within conditioned medium from patient-derived HFC and LFC region glioma cells (Extended Data Fig. 7a). Two-dimensional gel electrophoresis separating HFC and LFC proteins by size and charge produced up- and down- regulated candidate proteins known for their established role in neural signaling and tumorigenesis. Clusterin (CLU) was identified as an additional synaptogenic protein of interest with 1.4 upregulation in HFC compared with LFC samples (Extended Data Fig. 7b, c).

We co-cultured high-grade glioma cells from HFC and LFC tumor regions with mouse hippocampal neurons to test the effects of HFC and LFC patient-derived glioma cells on synaptic connectivity of neurons. In these 2D neuron-glioma co-cultures^3^, we established that neurofilament labels only neurons and that nestin labels only glioma cells and is not expressed in hippocampal neurons after 14 days in culture (Extended Data Fig. 8). We then quantified density of postsynaptic puncta (marked by the postsynaptic marker Homer-1) in HFC and LFC co-cultures. This quantification demonstrated increased postsynaptic Homer-1-positive puncta in glioma and neuronal processes in HFC co-cultures compared with LFC cultures (Fig. 2i), further indicating a role for HFC glioma cells in synaptogenesis.

### Neurons promote glioma circuit integration

Gliomas exhibit intratumoral heterogeneity with subpopulations of cancer cells assuming particular roles^18, 19^. Neuronal activity drives glioma growth and progression through activity-regulated paracrine signaling, direct neuron to glioma synaptic transmission and through action potential-dependent potassium-evoked currents^1–4^; each of these distinct forms of neuron-glioma interactions may be enriched in distinct subpopulations of glioma cells, such as enrichment of neuron-to-glioma synapses in the OPC-like glioma cells and activity-dependent potassium-evoked currents in more astrocyte-like glioma cells^3^. Similarly, preclinical studies have demonstrated a subpopulation of astrocyte-like glioma cells with synaptogenic properties that influence neuronal excitability and promote glioma-associated seizures^7, 28^. The human data presented above demonstrates geographic heterogeneity with respect to functional integration of the cancer with normal brain circuity and suggests that within intratumoral regions of high functional connectivity, a tumor subpopulation with synaptogenic properties exists. We next sought to further investigate functional distinctions between malignant subpopulations isolated form HFC and LFC regions by testing neuron-glioma interactions in neuronal organoid and in vivo settings. We co-cultured HFC and LFC glioma cells with GFP-labelled human neuron organoids, generated from an iPSC cell line integrated with a doxycycline-inducible human *NGN2* transgene to drive neuronal differentiation^32^. Quantification of postsynaptic Homer-1 in induced neuron (iN) organoids revealed a relative increase in postsynaptic puncta density when co-cultured with HFC glioma cells compared to LFC glioma cells (P = 0.0006) (Fig. 3a). Live cell imaging of neuronal organoids co-cultured with HFC and LFC glioma cells revealed that HFC glioma cultures exhibit prominent neuronal tropism and integrate extensively in the organoids, while LFC glioma cells displayed minimal integration with neuron organoids (Fig. 3b, Supplementary Videos 1, 2). Strikingly, exogenous administration of TSP-1 to iN-LFC co-culture reversed this phenotype and promoted robust glioma integration into the neuronal organoid (Fig. 3b, Supplementary Video 3), further implicating TSP-1 in neuron-glioma interactions.

**Fig. 3.**
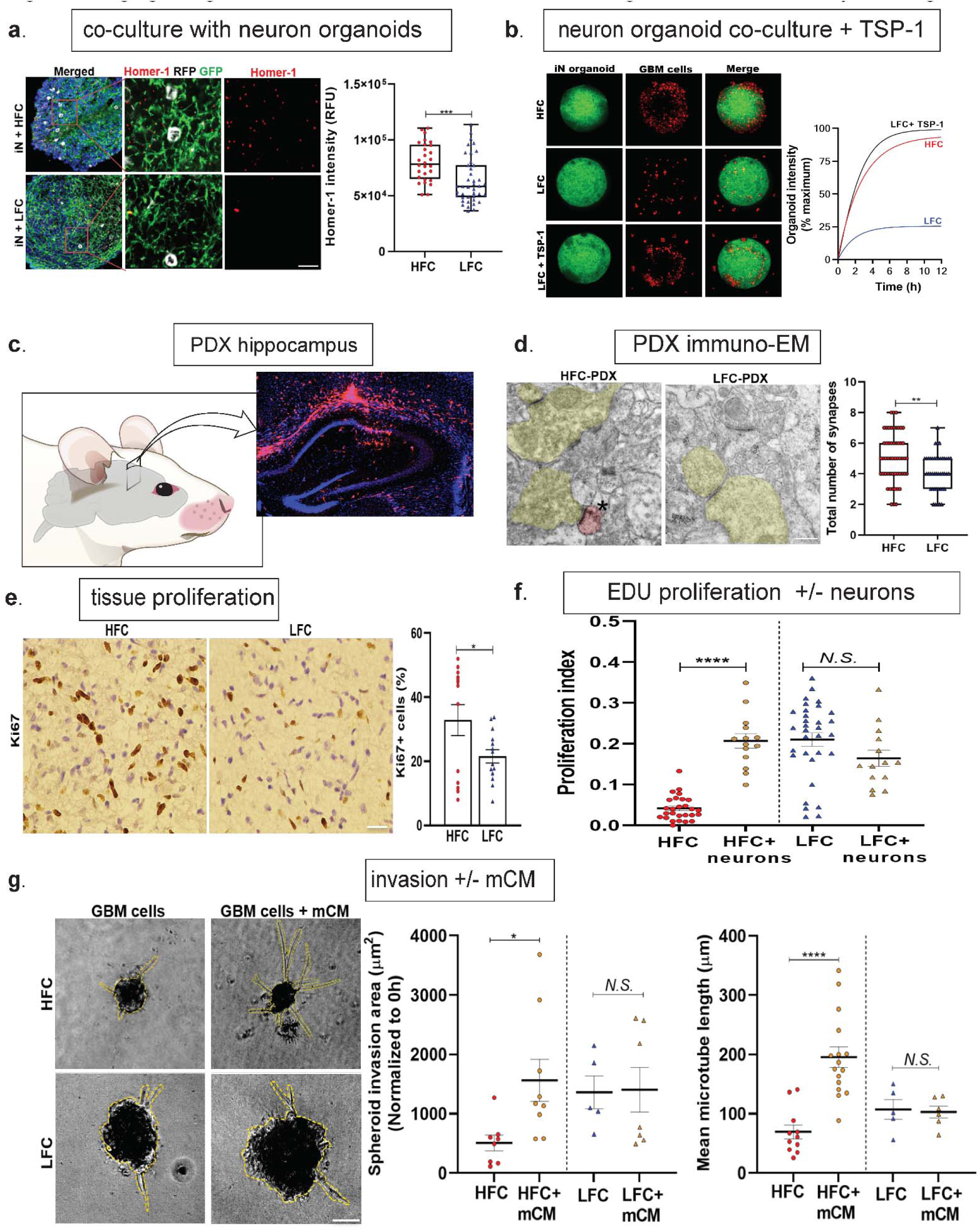
High-grade gliomas exhibit bidirectional interactions with high functional connectivity brain regions. a, Neuron organoids (GFP-labeled) were generated from an iPSC cell line integrated with doxycycline-inducible human NGN2 transgene and co-cultured with RFP labeled HFC and LFC cells (pseudo-colored white) for two weeks. Quantification of postsynaptic Homer-1 fluorescence intensity in 2-week induced neuron (iN) organoid sections expressed as relative fluorescence units (RFU; HFC vs. LFC: 80,162.21 ± 3,256.31 vs. 63,317.59 ± 3,185.48; n=1/group, n=2 organoids/group, n= 30-40 cells/organoid section analyzed). *P* = 0.0006. Scale bar, 10 µm. b, TSP-1 rescue of iN organoids in co-culture with HFC and LFC cells for 12 h. Quantification of glioblastoma (GBM) cell integration shown as percent of maximum for each condition from 0 to 100 (HFC vs. LFC vs. LFC+TSP-1: 68.93 ± 12.70 vs.19.98 ± 3.52 vs.75.16 ± 13.87; n =1/group). Scale bar, 300 µm. c, Representative micrograph showing RFP-labeled glioma cells xenografted into CA1 region of the mouse hippocampus. Scale bar, 500 µm. d, Immuno-electron microscopy of patient-derived HFC or LFC cells xenografted into the mouse hippocampus. Asterisk denotes immuno-gold particles labeling of RFP. Postsynaptic density in RFP+ tumor cells (pseudo-colored red), synaptic cleft, and clustered synaptic vesicles in apposing presynaptic neuron (pseudo-colored yellow) identify synapses. Quantification of total number of synapses (HFC vs. LFC: 5.06 ± 0.22 vs. 4.12 ± 0.19) in HFC/LFC xenografts (n = 4 mice per group). (*P* = 0.0019). Scale bar, 1000 nm. e, Representative immunohistochemistry images of the Ki67 proliferative marker staining in primary patient-derived tissues demonstrate increased protein expression in HFC samples (HFC vs. LFC: 32.92 ± 4.86% vs. 21.61 ± 2.11%, n= 4-5 images/sample, n= 4-5/group). *P* = 0.04. Scale bar, 50 µm. f, Primary patient glioblastoma (GBM) cells from high connectivity regions illustrate marked increase in proliferative index when co-cultured with mouse hippocampal neurons (HFC only vs. HFC + neurons vs. LFC vs. LFC + neurons: 0.04 ± 0.01 vs. 0.21 ± 0.02 vs. 0.21 ± 0.02 vs. 0.16 ± 0.02, n= 5-7 images/sample, n= 3-4/group). g, 3D spheroid invasion assay showing representative micrographs imaged 24 h after addition of invasion matrix. Analysis includes quantification of mean spheroid invasion area normalized for each sample to the initial (0 h) spheroid area (HFC vs. HFC + mCM vs. LFC vs. LFC + mCM: 507.96 ± 132.80 µm^2^ vs. 1,562.94 ± 353.95 µm^2^ vs. 1,361.38 ± 277.34 µm^2^ vs. 1,404.65 ± 376.94 µm^2^) and mean microtube length/spheroid (HFC vs. HFC + mCM vs. LFC vs. LFC + mCM: 69.45 ± 11.58 µm vs.195.22 ± 16.69 µm vs. 107.14 ± 16.64 µm vs. 102.95 ± 10.22 µm, n= 5-10 spheroids/sample, n= 1/group); n= 5-10 spheroids/sample, n= 1/group). Scale bar, 200 µm. Data presented as mean ± s.e.m (a-b, d-g). *P* values determined by two-tailed Student’s t-test. \**P* < 0.05. \*\**P* < 0.01. \*\*\**P* < 0.001. \*\*\*\**P* < 0.0001.

We then explored structural synapses in HFC glioma cell-infiltrated brain. RFP-labelled HFC or LFC glioma cells were stereotactically xenografted into the CA1 region of the mouse hippocampus^3^ (Fig. 3c). Following a period of engraftment and growth, immuno-electron microscopy analysis identified neuron-to-neuron and neuron-to-glioma synapses as previously reported^3, 4^, with RFP+ glioma cells on the postsynaptic side of synaptic structures with vesicle containing presynaptic boutons of non-RFP+ axons, a synaptic cleft, and clear post synaptic density on the RFP+ glioma process (Fig. 3d). HFC xenografts exhibited a greater number of synapses overall (neuron-to-neuron and neuron-to-glioma) compared to LFC xenografts, further demonstrating a greater synaptogenic potential of glioma cells isolated from HFC patient tumor regions.

Neurons promote glioma cell proliferation^1–4^ and we hypothesized that HFC cells may represent a cellular subpopulation within glioblastomas that are differentially regulated by neuronal factors. We found that primary patient biopsies from HFC and LFC regions demonstrated increased Ki-67 proliferative marker staining within HFC regions (32.9% Ki-67 expression in HFC samples compared with 21.6% LFC samples, *P* = 0.04) (Fig. 3e). To test whether HFC cells differentially proliferate in response to neuronal factors when compared with LFC primary patient cultures, patient-derived HFC and LFC cells cultured alone or in co-culture with mouse hippocampal neurons were treated with EdU overnight. HFC glioma cells exhibit a 5-fold increase in proliferation when cultured with neurons (from 4% to 21% EdU+ cells). In contrast, LFC glioma *in vitro* cell proliferation index (determined as the fraction of DAPI cells co-expressing EdU) is similar with and without hippocampal neurons in vitro (Fig. 3f, Extended Data Fig. 9a, b).

Given the neuronal tropism exhibited by HFC glioma cells together with the concept that neural network integration requires invasion of brain parenchyma in order to reach and co-localize with neuronal elements, we next tested the effects of neuronal conditioned medium on invasion of HFC and LFC glioma cells using a 3D spheroid invasion assay. LFC glioma cells demonstrated no differences in spheroid volume in the presence or absence of neuronal conditioned medium, however HFC glioma cells exhibited increased spheroid invasion area in response to neuronal conditioned medium. In addition to increased invasion area, HFC glioma cells extended long processes representing tumor microtubes^33^ in response to neuronal conditioned medium. Tumor microtubes connect glioma cells in a gap junction-coupled network ^4, 33, 34^ through which neuronal activity-induced currents are amplified^3^. Quantifying the change in mean spheroid volume as a measure of invasion as well as microtube length, we found that neuronal conditioned medium increased both invasion and microtube length in HFC but not LFC cultures (Fig. 3g). Concordantly, the invasive marker MET was increased within HFC samples compared with LFC (Extended Data Fig. 10). Taken together, these results suggest that functionally connected intratumoral regions are enriched for a tumor cell population that is differentially responsive to neuronal signals and exhibits proliferative, invasive, and integrative phenotype in the neuronal microenvironment.

### Glioma functional connectivity shortens survival

We next explored the effects of high functional connectivity within gliomas on survival and cognition. First, we tested the hypothesis that gliomas exhibiting increased functional connectivity may be more aggressive, given the robust influence of neuronal activity on tumor progression^1–3^. We performed survival studies of mice orthotopically xenografted with patient-derived HFC or LFC glioma cells. Mice bearing HFC tumors exhibited shorter survival compared to LFC xenografted mice (Fig. 4a). To investigate patient outcomes, we performed a human survival analysis of patients with molecularly uniform newly diagnosed IDH-WT glioblastoma. After controlling for known correlates of survival (age, tumor volume, and extent of tumor resection)^35^, neural oscillations and functional connectivity were measured within tumor-infiltrated brain using MEG (Extended Table 4). Subjects were classified by the presence or absence of HFC voxels within the tumor boundary. Kaplan-Meier survival analysis illustrates 71-week overall survival for patients with HFC voxels as compared to 123- week overall survival for participants without HFC voxels, illustrating a striking inverse relationship between survival and functional connectivity of the tumor (mean follow-up months 50.5 months) (*P* = 0.04) (Fig. 4b). We hypothesized that, beyond survival, intratumoral functional connectivity may also influence cognition. We therefore performed visual picture naming and auditory stimulus naming tests identical to intraoperative physiology testing (Fig. 1e) in our cohort of patients with dominant hemisphere IDH-WT glioblastoma (Fig. 4c). Picture naming and auditory stimulus naming language tasks are sensitive predictors of aphasia in clinical populations^36, 37^. Linear regression of the number of HFC voxels within tumor with language task performance demonstrated an inverse relationship between language cognitive performance and tumor functional connectivity (Fig. 4d). Together, these findings suggest that functional integration of glioblastoma into neural circuits negatively influences cognition and survival.

**Fig. 4.**
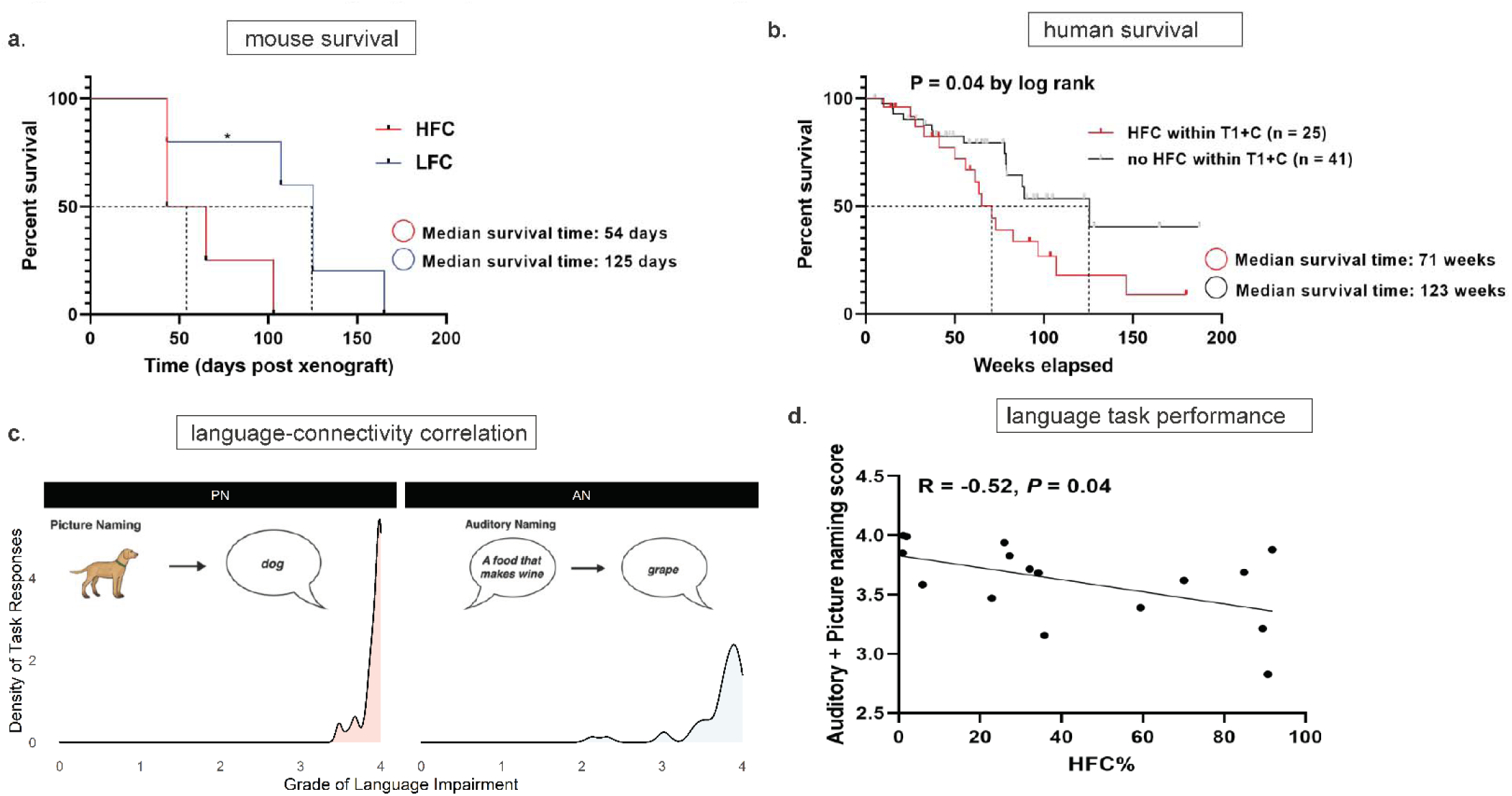
Intratumoral connectivity in HGG patients correlates with survival and cognition. a, Kaplan-Meier survival curves of mice bearing HFC or LFC xenografts; n = 4-5 mice per group. \**P* < 0.05. *P*-value determined by two-tailed log rank analysis. b, Kaplan-Meier human survival analysis illustrates 71-week overall survival for patients with HFC voxels as determined by contrast-enhanced T1-weighted images as compared to 123-weeks for participants without HFC voxels (mean follow-up months 50.5, range 4.9-155.9 months). c, Picture and auditory naming language task performance across the study population. d, Linear regression of HFC% (calculated by dividing the number of intrahemispheric HFC voxel counts by the total number of intrahemispheric functional connectivity [HFC+LFC] counts) with baseline auditory and picture naming scores (n= 16). \**P* < 0.05, two-tailed log rank analysis (a, b).

## Discussion

High-grade glioma integration into neural networks is manifested by bidirectional interactions whereby neuronal activity increases glioma growth^1–3^ and gliomas increase neuronal excitability^5–8^. To understand whether glioma-neuronal interactions influence neural circuit dynamics, we used short-range electrocorticography of tumor-infiltrated human brain regions to demonstrate task-specific language activation of tumor-infiltrated cortex as well as remodeling of language circuits such that language tasks activate the entire glioma-infiltrated cortex. We demonstrated in humans with glioblastoma, using visual and auditory word retrieval as an illustrative case, that high-grade gliomas remodel functional circuits and that distinct intratumoral regions maintain functional connectivity through a subpopulation of *TSP-1* expressing malignant cells (HFC glioma cells). In addition, we show that this molecularly distinct glioma subpopulation is differentially responsive to neuronal signals, exhibiting a synaptogenic, proliferative, invasive, and integrative signature. Past work has demonstrated that neuronal activity promotes glioma proliferation through paracrine and synaptic signaling^1–4^, and we have now shown that patients with high-grade gliomas exhibiting intratumoral functional connectivity experience shorter overall survival when compared with patients without high functional connectivity.

The neuronal microenvironment has emerged as a crucial regulator of glioma growth. Both paracrine signaling as well as connectivity remodeling may contribute to network level changes in patients impacting both cognition and survival. In patients, the role of network dynamics on survival and cognition remains poorly understood. In fact, using a heterogenous population of patients with both IDH-wild type and mutant WHO grade III and IV gliomas, some have suggested that functional connectivity improves overall survival^38^ and how glioma-network interactions influences cognition remains unanswered; such prior work has been confounded by functional connectivity methods heavily influenced by the presence of tumor vascularity, limited spatial resolution, in addition to a heterogenous patient cohort. A better understanding of the cross-talk between neurons and gliomas as well as how functional integration impacts clinical outcomes may open the door to a range of pharmacological and neuromodulation therapeutic strategies focused on improving cognitive outcomes as well as survival.

## Methods

We began by studying short-range circuit dynamics in a subset of 14 patients with dominant hemisphere gliomas infiltrating speech production areas of the inferior frontal lobe using electrocorticography (ECoG) in the intraoperative setting (Fig. 1a). We then focused molecular studies on patients with surgically treated IDH-wild type (IDH WT) glioblastoma, performed extra- operative language assessments and imaginary coherence as a long-range measure of functional connectivity using magnetoencephalography (MEG) (Fig. 1a-b). This allowed us to import functional connectivity data into the operating room where we performed site-specific tissue biopsies of human glioma from regions with differing measures of functional connectivity for in vivo and in vitro cell biology experiments including primary patient cultures (*n* = 19 patients) and multimodal tissue profiling, including microscopy, sequencing, proteomics, and patient-derived tumor xenografting (Fig. 1c). This layered approach combining clinical variables, cognition assessments, human and animal models of network dynamics, in addition to cell biology served as a platform to study the clinical implications of glioma-neuron interactions (Fig. 1d, Extended Data Table 1).

### Human Electrocorticography (ECoG) and Data Analyses

Intra-operative photographs with and without subdural electrodes present were used to localize each electrode contact. Images were registered using landmarks from gyral anatomy and vascular arrangement to preoperative T1- and T2-weighted MRI scans. Tumor boundaries were localized on MRI scans and electrodes within 10 mm of necrotic tumor core tissue were identified as ‘tumor’ contacts. Electrodes overlying the hypointense core of the tumor were identified as “core”, electrodes extending from the contrast enhancing rim extending to the edge of FLAIR were considered “infiltrative margin”, and electrodes completely outside of any T1 post gadolinium or FLAIR signal were considered “non-tumor or healthy”.

Electrocorticographic (ECoG) signals were acquired during a period after stopping the administration of anesthetics and the patient was judged to be alert and awake^39^. Intraoperative tasks consisted of naming pictorial representations of common objects and animals (Picture Naming, PN) and naming common objects and animals via auditory descriptions (Auditory Naming, AN)^40^. Post-operative videos were re-analyzed to ensure all data was collected and correct responses only included for analysis. Recordings were acquired at 4800 Hz and down-sampled to 1200 Hz during the initial stages of processing. Channels with excessive noise artifacts were visually identified and removed. Following the rejection of artifactual channels, data were referenced to a common average, high-pass filtered at 0.1 Hz to remove slow drift artifacts, and bandpass filtered between 70–110 Hz using a 300-Order FIR filter to focus the analyses on the high-gamma band range, which is strongly related to local mean population spiking rates. High-gamma band power (HGp) was then calculated using the square of the Hilbert transform on the filtered data. HGp was then averaged across the resting-state time-series, yielding a single measure of neural responsivity for each electrode contact. HGp was then averaged across patients during the task response period, yielding a single measure of neuronal responsivity for each channel. HGp levels were then compared between tumor and non-tumor channels using independent samples t-tests separately for each patient.

### Magnetoencephalography (MEG) recordings and data analysis

MEG recordings were performed according to an established protocol^24, 41^. Briefly, study participants had continuous resting state MEG recorded with a 275-channel whole- head CTF Omega 2000 system (CTF Systems, Inc., Coquitlam, BC, Canada) using a sampling rate of 1200Hz. During resting state recordings participants are awake with eyes closed. Surface landmarks were co-registered to structural magnetic resonance (MR) images to generate the head shape. An artifact-free 1-minute epoch was selected for further analysis if the patient’s head movement did not exceed 0.5 cm. This artifact-free, 1-minute epoch was then analyzed using the NUTMEG software suite (UCSF Biomagnetic Imaging Laboratory) to reconstruct whole-brain oscillatory activity from MEG sensors so as to construct functional connectivity (imaginary coherence [IC]) metrics^42–44^. Spatially normalized structural MR images were used to overlay a volume-of-interest projection (grid size = 8 mm; approximately 3000 voxels/subject) such that each voxel contained the entire time series of activity for that location derived by all the MEG sensor recordings. The time series within each voxel was then bandpass filtered for the alpha band (8-12 Hz) and reconstructed in source space using a minimum-variance adaptive spatial filtering technique^42, 45^. The alpha frequency band was selected because it was the most consistently identified peak in the power spectra from this sampling window in our patient series. Functional connectivity estimates were calculated using IC, a technique known to reduce overestimation biases in MEG data generated from common references, cross-talk, and volume conduction^22, 23^.

### Functional connectivity map

The functional connectivity of an individual voxel was derived by the mean IC between the index voxel and the rest of the brain, referenced to its contralesional pair^41^. In comparison to contralesional voxels, we used a two-tailed t-test to test the null hypothesis that the Z-transformed connectivity IC between the index voxel and nontumor voxel is equal to the mean of the Z-transformed connectivity between all contralateral voxels and the same set of voxels. The resultant functional connectivity values were separated into tertiles: upper tertile (high functional connectivity [HFC]) and lower tertile (low connectivity [LFC]). Functional connectivity maps were created by projecting connectivity data onto each individual patient’s preoperative structural MR images. Functional connectivity maps were created and projected onto structural MR images for each subject and imported into the operating room neuro-navigation console. Stereotactic site-directed biopsies from HFC (upper tertile) and LFC (lower tertile) intratumoral regions were taken and X, Y, Z coordinates determined using Brainlab neuro-navigation.

### Isolation and culture of primary patient-derived glioblastoma cells

For all human tissue studies, informed consent was obtained, and tissue was used in accordance with the University of California, San Francisco (UCSF) institutional review board (IRB) for human research. Tumor tissues with high (HFC) and low (LFC) functional connectivity sampled during surgery based on preoperative MEG were subjected to quality control by a certified neuropathologist and subsequently used to generate primary patient-derived cultures. Patient-matched samples were acquired from site directed HFC and LFC intratumoral regions from the same patient. Intratumoral HFC and LFC tissues were dissociated both mechanically and enzymatically and then passed through a 40 µm filter to remove debris. The filtered cell suspension was then treated with ACK lysis buffer (Invitrogen) to remove red blood cells and subsequently cultured as free-floating neurospheres in a defined, serum-free medium designated ‘tumor sphere culture media’ (TSM), consisting of Dulbecco’s Modified Eagle’s Medium (DMEM-F12; Invitrogen), B27 (Invitrogen), N2 (Invitrogen), human-EGF (20 ng/ml; Peprotech), human-FGF (20 ng/ml; Peprotech). In addition, Normocin (InvivoGen) was added to the cell culture medium in combination with Penicillin/Streptomycin (Invitrogen) to prevent mycoplasma, bacterial and fungal contaminations. Cell cultures were routinely tested for Mycoplasma (PCR Mycoplasma Test Kit I/C, PromoCell, Heidelberg, Germany) and no positive results were obtained (Extended Data Fig. 11).

### Bulk RNA-sequencing and analysis

RNA was isolated from HFC (n = 3) and LFC (n = 4) tumor samples using the RNeasy Plus Universal Mini Kit (QIAGEN, Valencia, CA) and RNA quality was confirmed using an Advanced Analytical Fragment Analyzer. RNA sequencing libraries were generated using the TruSeq Stranded RNA Library Prep Kit v2 (RS-122- 2001, Illumina, San Diego, CA) and 100 bp paired-end reads were sequenced on an Illumina HiSeq 2500 to at least 26 million reads per sample at the Functional Genomics Core Facility at UCSF. Quality control of FASTQ files was performed with FASTQC (http://www.bioinformatics.babraham.ac.uk/projects/fastqc/). Reads were trimmed with Trimmomatic^46^ to remove leading and trailing bases with quality scores below 20 as well as any bases that did not have an average quality score of 20 within a sliding window of 4 bases. Any reads shorter than 72 bases after trimming were removed. Reads were subsequently mapped to the human reference genome GRCh38^47^ using HISAT2^48^ version 2.1.0 with default parameters. For differential expression analysis, we extracted exon level count data from the mapped HISAT2 output using featureCounts^49^. Differentially expression analysis was done with DESeq2^50^, using the ‘apeglm’ parameter^51^ to accurately calculate log fold changes and setting a false discovery rate of 0.05. Differentially expressed genes were identified as those with log fold changes greater than 1 and an adjusted *P* value less than 0.05. Unsupervised gene expression principal component analysis (PCA) and volcano plots of IDH-WT glioblastoma (Extended Data Fig. 3a, b) revealed 144 differentially expressed genes between HFC and LFC tumor regions including 40 genes involved in nervous system development (Extended Table 2).

## Single-cell sequencing

### Single-cell suspension generation

Fresh tumor samples were acquired from the operating room and transported to the laboratory space in PBS and on ice. Tumor tissue was minced with #10 scalpels (Integra LifeSciences, Plainsboro Township, NJ) and then digested in papain (Worthington Biochemical LK003178, Lakewood, NJ) for 45 minutes at 37°C. Digested tumor tissue was then incubated in red blood cell lysis buffer (eBioscience 00-4300-54, San Diego, CA) for 10 minutes at room temperature. Finally, samples were sequentially filtered through 70- and 40-micron filters to generate a single-cell suspension.

### Single-cell sequencing and analysis

Single-cell suspensions of 3 patient-matched HFC and LFC tumor tissues were generated as described above and processed for single-cell RNA-seq using the Chromium Next GEM Single Cell 3’ GEM, Library & Gel Bead Kit v3.1 on a 10x Chromium controller (10x Genomics, Pleasanton, CA) using the manufacturer recommended default protocol and settings, at a target cell recovery of 5,000 cells per sample. 100 base pair paired-end reads were sequenced on an Illumina NovaSeq 6000 at the Center for Advanced Technology at the University of California San Francisco, and the resulting FASTQ files were processed using the CellRanger analysis suite version 3.0.2 (https://github.com/10XGenomics/cellranger) for alignment to the hg38 reference genome, identification of empty droplets, and determination of the count threshold for further analysis. A cell quality filter of greater than 500 features but fewer than 10,000 features per cell, and less than 20% of read counts attributed to mitochondrial genes, was used. Single cell UMI count data were preprocessed in Seurat 3.0.1^52, 53^ using the sctransform workflow^54^, with scaling based on regression of UMI count and percentage of reads attributed to mitochondrial genes per cell. Dimensionality reduction was performed using principal component analysis and then principal component loadings were corrected for batch effects using Harmony^55^. Uniform Manifold Approximation and Projection (UMAP) was performed on the reduced data with a minimum distance metric of 0.4 and Louvain clustering was performed using a resolution of 0.2. Marker selection was performed in Seurat using a minimum difference in the fraction of detection of 0.5 and a minimum log-fold change of 0.5. We assessed the single cell transcriptome from 6,666 HFC region cells and 7,065 LFC region cells (Extended Data Table 3).

### ELISA

Peripheral blood samples from newly diagnosed glioblastoma patients (n = 16) were collected and allowed to clot for 30 minutes at room temperature before centrifugation for 15 minutes at 1000 x g. The serum was stored at −80°C until analysis. TSP-1 level was determined using the Quantikine immunosorbent assay kits (DTSP10) as per the manufacturer’s instructions (R&D Systems, Minneapolis, MN, USA).

### Orthotopic xenografting for neuronal circuit integration experiments

For all xenograft studies, NSG mice (NOD-SCID-IL2R gamma chain-deficient, The Jackson Laboratory) were used. Male and female mice were used equally. For immuno-electron microscopy experiment, a single-cell suspension from cultured neurospheres of HFC and LFC (N= 2 each) labeled with red fluorescent protein (RFP) were prepared in sterile DMEM immediately before the xenograft procedure. Mice (n = 8; 2 biological replicates/patient line) at postnatal day (P) 28–30 were anaesthetized with 1–4% isoflurane and placed in a stereotactic apparatus. The cranium was exposed via midline incision under aseptic conditions. Approximately 50,000 cells in 2 µl sterile PBS were stereotactically implanted into the CA1 region of the hippocampus through a 31-gauge burr hole, using a digital pump at infusion rate of 0.4 µl min−1 and 31-gauge Hamilton syringe. Stereotactic coordinates used were as follows: 1.5 mm lateral to midline, 1.8 mm posterior to bregma, −1.4 mm deep to cranial surface. At the completion of infusion, the syringe needle was allowed to remain in place for a minimum of 2 min, then manually withdrawn at a rate of 0.875 mm/min to minimize backflow of the injected cell suspension.

### Sample preparation and image acquisition for electron microscopy

Twelve weeks after xenografting, mice were euthanized by transcardial perfusion with Karnovsky’s fixative: 2% glutaraldehyde (EMS 16000) and 4% paraformaldehyde (EMS 15700) in 0.1M sodium cacodylate (EMS 12300), pH 7.4. Transmission electron microscopy (TEM) was performed in the tumor mass within the CA1 region of the hippocampus for all xenograft analysis. The samples were then post-fixed in 1% osmium tetroxide (EMS 19100) for 1 h at 4°C, washed 3 times with ultrafiltered water, then en bloc stained for overnight at 4 °C. Samples were dehydrated in graded ethanol (50%, 75% and 95%) for 15 min each at 4 °C; the samples were then allowed to equilibrate to room temperature and were rinsed in 100% ethanol twice, followed by acetonitrile for 15 min. Samples were infiltrated with EMbed-812 resin (EMS 14120) mixed 1:1 with acetonitrile for 2 h followed by 2:1 EMbed-812:acetonitrile overnight. The samples were then placed into EMbed-812 for 2 h, then placed into TAAB capsules filled with fresh resin, which were then placed into a 65L overnight. Sections were taken between 40 and 60 nm on a Leica Ultracut S (Leica) and mounted on 100-mesh Ni grids (EMS FCF100-Ni). For immunohistochemistry, microetching was done with 10% periodic acid and eluting of osmium with 10% sodium metaperiodate for 15 min at room temperature on parafilm. Grids were rinsed with water three times, followed by 0.5 M glycine quench, and then incubated in blocking solution (0.5% BSA, 0.5% ovalbumin in PBST) at room temperature for 20 min. Primary goat anti-RFP (1: 300, ABIN6254205) was diluted in the same blocking solution and incubated overnight at 4 °C. The next day, grids were rinsed in PBS three times, and incubated in secondary antibody (1:10 10-nm gold-conjugated IgG TED Pella 15796) for one hour at room temperature and rinsed with PBST followed by water. For each staining set, samples that did not contain any RFP-expressing cells were stained simultaneously to control for any non-specific binding. Grids were contrast stained for 30 s in 3.5% uranyl acetate in 50% acetone followed by staining in 0.2% lead citrate for 90 s. Samples were imaged using a JEOL JEM-1400 TEM at 120 kV and images were collected using a Gatan Orius digital camera.

### Electron microscopy data analysis

Sections from the xenografted hippocampi of mice were imaged as above using TEM imaging. Here, 101 sections of HFC xenografts across 4 mice and 104 sections of LFC xenografts across 3 mice were analyzed. Electron microscopy images were taken at 6,000× with a field of view of 15.75 μ . Glioma cells were counted and analyzed after unequivocal identification of a cluster of immunogold particle labelling with fifteen or more particles. The total number of synapses, including neuron-to-neuron and neuron-to-glioma synapses (identified by: (1) presence of synaptic vesicle clusters; (2) visually apparent synaptic cleft; and (3) identification of clear postsynaptic density in the glioma cell) were counted.

### Orthotopic xenografting for mouse survival experiments

For survival studies, five-week old athymic mice, purchased from Harlan (Indianapolis, IN) were used. A single-cell suspension from cultured neurospheres of HFC and LFC (n= 2-3 per group) were prepared in sterile DMEM immediately before the xenograft procedure. Mice (n = 4-5 mice per group) were anaesthetized with 1–4% isoflurane and placed in a stereotactic apparatus. The cranium was exposed via midline incision under aseptic conditions. Approximately 40,000 cells in 2 µl sterile PBS were stereotactically implanted into the prefrontal cortex through a 31-gauge burr hole, using a digital pump at infusion rate of 0.4 µl min−1 and 31-gauge Hamilton syringe. Stereotactic coordinates relative to bregma in millimeters were 1.7 AP, 0.3 ML, and −2.5 DV. At the completion of infusion, the syringe needle was allowed to remain in place for a minimum of 2 min, then manually withdrawn at a rate of 0.875 mm/min to minimize backflow of the injected cell suspension. For survival studies, morbidity criteria used were either: reduction of weight by 15% initial weight, or clinical signs such as hunched posture, lethargy, or persistent decumbency. Kaplan-Meier survival analysis using log rank testing was performed to determine statistical significance.

### Glioma-mouse hippocampal neuron co-culture

Glioma cells were plated on poly-d-lysine and laminin coated coverslips (Neuvitro) at a density of 10,000 cells/well in 24-well plate. Approximately 24 h later, 40,000 embryonic mouse hippocampal neurons (Gibco) were seeded on top of the glioma cells and maintained with serum-free Neurobasal medium supplemented with B27, gentamicin and Glutamax (Gibco). After 2 weeks of co-culture, cells were fixed with 4% paraformaldehyde (PFA) for 30 min at 4°C and incubated in blocking solution (5% normal donkey and goat serum, 0.25% Triton X-100 in PBS) at room temperature for 1 h. Next, they were treated with primary antibodies diluted in the blocking solutions overnight at 4°C. The following antibodies were used: rabbit anti-homer-1 (1:500, Pierce) mouse anti-nestin (1:500, Abcam), chicken anti-neurofilament (M+H; 1:1,000; Aves Labs). Coverslips were then rinsed three times in PBS and incubated in secondary antibody solution (Alexa 488 goat anti-chicken IgG; Alexa 568 goat anti-mouse IgG, and Alexa 647 goat anti-rabbit IgG all used at 1:500 (Invitrogen) in antibody diluent solution for 1 h at room temperature. Coverslips were rinsed three times in PBS and then mounted with VECTA antifade mounting media with 4, 6-diamidino-2-phenylindole dihydrochloride (DAPI, Vector Laboratories).

### Confocal imaging and quantification of Homer-1 staining

Images were captured at 1024 x 1024 resolution using a 10X objective on a Nikon C2 confocal microscope. Area of imaging was selected solely on basis of the nestin 568 channel for glioma cells, and hence the experimenter was blinded to the Homer-1 staining. The confocal microscope settings for the Homer Alexa647 channel were held constant across all the samples used for the experiment. Collected images were then imported to Imaris software (Imaris 9.2.1, Bitplane) and the spots detection function of Imaris was applied to the Cy5 (Alexa647) channel. Square ROIs of 285 µm^2^ was set around each glioma cell in proximity with neurofilament-positive neurons and the selection area was then kept constant across all images. Homer-1 puncta fluorescence was quantified from each ROI using the spot detection function of Imaris, with automatic intensity max spot detection thresholds and a spot diameter of 1.0 µm. Quantification of Homer-1 puncta density was calculated by dividing the number of puncta measured with the area of the image field.

### Immunohistochemistry and Immunofluorescence

After rehydration, 5.0 μm paraffin-embedded sections were subjected to antigen retrieval followed by blocking and primary antibody incubation overnight at 4°C. The following primary antibodies were used: rabbit anti-synapsin 1 (1:1000, EMD Millipore), mouse anti-PSD95 (1:100, UC Davis), mouse anti-nestin (1:500, Abcam), mouse anti-neurofilament (M+H; 1:1000, Novus Biologicals), mouse anti-TSP-1 (1:50, Invitrogen), rabbit anti-MET (1:100, Abcam), rabbit anti-Ki67 (1:100, Abcam). We used species-specific secondary antibodies tagged with Alexa Fluor 488, 568, or 647 (1:1,000, Invitrogen) for immunofluorescence. After DAPI nuclear counter staining (Vector Laboratories, 1:1,000), coverslips were mounted with Fluoromount-G mounting medium (SouthernBiotech). DAB horseradish peroxidase (HRP) (Vector Laboratories) was used for chemical colorimetric detection following species-specific, HRP-conjugated secondary antibody labelling. The number of synapsin-1 and PSD95 puncta was quantified using spots (with automatic intensity max spot detection thresholds and a spot diameter of 1.0 µm) detection function of Imaris. The ratio of pre-and postsynaptic puncta was calculated by dividing the total number of synapsin-1 or PSD95 puncta on neurofilament-positive neurons to the total number of cells stained with DAPI in 135 x 135-μ

### EdU assay

The 5-ethynyl-2’-deoxyuridine (EdU) incorporation assay was performed with an EdU assay kit (Invitrogen) according to the manufacturer’s instructions. Patient-derived HFC/LFC glioma cells were seeded on poly-D-lysine and laminin coated coverslips at 10,000 cells per well of a 24 well plate. After 24 h of seeding, embryonic mouse hippocampal neurons were added to the glioma cells at 40,000 cells per well for the glioma-neuron co-culture group. After 72 h, glioma cells alone or in coculture with neurons were treated with 20 μM EdU overnight at 37 °C. Subsequently, the cells were fixed with 4% paraformaldehyde and stained using the Click-iT EdU kit protocol. Proliferation index was then determined by quantifying the fraction of EdU labeled cells/DAPI labeled cells using confocal microscopy at 10x magnification.

### Induced neuron organoid and glioma co-culture

Induced neuron organoids were generated from WTC11 human induced pluripotent stem cells (iPSCs) clone integrated by human NGN2 transgene induction as described previously^32, 56^. Briefly, iNeuron organoids were generated by transgenic hiPSC WTC11 line by NGN2 induction via addition of 2ug/ml doxycycline in the 1:1 mixture of Neurobasal^TM^ and BrainPhys^TM^ neural medium containing 1% B-27 supplement, 0.5% GlutaMAX, 0.2uM compound E, 10ng/ml BDNF, and 10ng/ml NT-3 for 10 days to induce neuronal differentiation. Following this, neuron maturation was triggered by feeding the organoids with approximately 8-month-old organoid conditioned medium derived from astrocytes. Astrocytes were differentiated from hiPSC WTC11 line and cultured in a medium consisting of DMEM/ F12 containing glutamax, sodium bicarbonate, sodium pyruvate, N-2 supplement, B-27 supplement (Gibco), 2ug/mL heparin, 10ng/ml EGF and 10ng/ml FGF2. Neuron organoids were characterized as postmitotic and stained for MAP2 and βIII-Tubulin to validate neuronal induction efficiency. After 14 days of neuronal differentiation, HFC/LFC glioma cells labeled with RFP were added to the neuron organoid culture at a ratio of 1:3. Prior to iNeuron induction, transgenic hiPSC WTC11 was transduced with GFP lentivirus. A Zeiss Cell Observer Spinning Disc Confocal microscope (Carl Zeiss AG, Oberkochen, Germany) fitted with a temperature and carbon dioxide-controlled chamber was used to record live interactions of glioma cells with neuron organoids. Organoids were imaged every 10 min for a 12 h period, starting at the time of co-culture initiation, using a 10× objective with 0.4 NA. To assess the effect of exogenous TSP-1 on the functional integration between glioma cells and neurons, human recombinant TSP-1 (R&D system Minneapolis, MN) was applied at a dose of 5 µg/ml to the LFC-neuron organoid co-culture. At the end of two weeks, organoids from HFC and LFC co-cultures were embedded in OCT and sectioned at 10 um thickness for Homer-1 immunofluorescence staining.

### 3D spheroid invasion assay

Glioma cell invasion was evaluated by performing invasion assay using the Cultrex 3D Spheroid Cell Invasion Assay Kit (Trevigen, Inc., Gaithersburg, MD) as per manufacturer’s protocol. Briefly, 3,000 cells were resuspended in 50 ul of spheroid formation matrix solution (prepared in culture media) in a round bottom 96-well plate. Spheroids were allowed to form for 72 h and images were taken at 10X magnification before addition of invasion matrix (0 h). Working on ice, 50 μl of invasion matrix was then added to each well and the plate was incubated at 37 °C. After 1 h of gel formation, for the wells to assess the effect of neuronal secreted factors on the invasiveness of the glioma cells. After 24 h of incubation at 37 °C, invasions were observed under microscope and images were taken at 10X. Microtube lengths as well as the area of each spheroid measured at 0 h (pre-invasion) and 24 h (post-invasion) were analyzed using Image J and the difference was used to calculate the total area of cell invasion.

### Proteomics analysis

The supernatants from one-month-old primary patient HFC and LFC cultures (n =3 each) were collected and sent to Applied Biomics (Hayward, CA) for two-dimensional gel electrophoresis (2-D DIGE). Differentially altered protein spots were further selected and identified by mass spectrometry (MALDI-TOF/TOF) using GPS Explorer software equipped with the MASCOT.

### Two-Dimensional Gel Electrophoresis

Two-dimensional gel electrophoresis (2-D DIGE) and subsequent Protein ID were performed by Applied Biomics, Inc (Hayward, CA).

#### Preparation of Samples and CyDye Labeling

Protein sample buffer was exchanged into 2-D cell lysis buffer (30 mM Tris-HCl, pH 8.8, containing 7 M urea, 2 M thiourea and 4% CHAPS). Protein concentration was measured using Bio-Rad protein assay method (Hercules, CA). For each sample, 30 μg of protein was mixed with 1.0 μl of diluted CyDye, and kept in the dark on ice for 30 min. The labeling reaction was stopped by adding 1.0 μ of 10 mM Lysine to each sample, and incubating in the dark on ice for an additional 15 min. The labeled samples were then mixed together. The 2X 2-D Sample buffer (8 M urea, 4% CHAPS, 20 mg/ml DTT, 2% pharmalytes and trace amount of bromophenol blue), 100 μl destreak solution and Rehydration buffer (7 M urea, 2 M thiourea, 4% CHAPS, 20 mg/mL DTT, 1% pharmalytes and trace amount of bromophenol blue) were added to the labeling mix to make the total volume of 250 μl for the 13-cm IPG strip.

#### IEF and SDS-PAGE

After loading the labeled samples, IEF (pH 3-10) was run following the protocol provided by GE Healthcare. Next, the IPG strips were incubated in the freshly made equilibration buffer-1 (50 mM Tris-HCl, pH 8.8, containing 6 M urea, 30% glycerol, 2% SDS, trace amount of bromophenol blue and 10 mg/mL DTT) for 15 min with gentle shaking. Then the strips were rinsed in the freshly made equilibration buffer-2 (50 mM Tris-HCl, pH 8.8, containing 6 M urea, 30% glycerol, 2% SDS, trace amount of bromophenol blue and 45 mg/mL Iodoacetamide) for 10 min with gentle shaking. Next, the IPG strips were rinsed in the SDS-gel running buffer before transferring into 12% SDS-gels. The SDS-gels were run at 15°C until the dye front ran out of the gels.

#### Image Scan and Data Analysis

Gel images were scanned immediately following the SDS-PAGE using Typhoon TRIO (GE Healthcare, Waukesha, WI). The scanned images were then analyzed by Image Quant software (version 6.0, GE Healthcare, Waukesha, WI), followed by quantitation analysis using DeCyder software (version 6.5, GE Healthcare, Waukesha, WI). The fold change of the protein expression levels was obtained from in-gel DeCyder analysis.

### Protein Identification by Mass Spectrometry

#### Spot Picking and Trypsin Digestion

The spots of interest were picked up by Ettan Spot Picker (Amersham BioSciences, Piscataway, NJ) based on the in-gel analysis and spot picking design by DeCyder software. The gel spots were washed a few times, then digested in-gel with modified porcine trypsin protease (Trypsin Gold, Promega, Madison, WI). The digested tryptic peptides were desalted by Zip-tip C18 (Millipore, Billerica, MA). Peptides were eluted from the Zip-tip with 0.5 μl of matrix solution (α-cyano-4-hydroxycinnamic acid 5 mg/mL in 50% acetonitrile, 0.1% trifluoroacetic acid, 25 mM ammonium bicarbonate) and spotted on the AB SCIEX MALDI plate (Opti-TOF 384 Well Insert, AB SCIEX, Framingham, MA).

#### Mass Spectrometry

MALDI-TOF MS and TOF/TOF tandem MS/MS were performed on an AB SCIEX TOF/TOF 5800 System (AB SCIEX, Framingham, MA). MALDI-TOF mass spectra were acquired in reflectron positive ion mode, averaging 4000 laser shots per spectrum. TOF/TOF tandem MS fragmentation spectra were acquired for each sample, averaging 4000 laser shots per fragmentation spectrum on each of the 10 most abundant ions present in each sample (excluding trypsin autolytic peptides and other known background ions).

### Database Search

Both the resulting peptide mass and the associated fragmentation spectra were submitted to GPS Explorer workstation equipped with MASCOT search engine (Matrix Science, Boston, CA) to search the Swiss-Prot database. Searches were performed without constraining protein molecular weight or isoelectric point, with variable carbamidomethylation of cysteine and oxidation of methionine residues, and with one missed cleavage also allowed in the search parameters. Candidates with either protein score C.I.% or Ion C.I.% greater than 95 were considered significant.

### Measurement of tumor volume and calculation of volumetric extent of resection

Pre-operative and post-operative tumor volumes were quantified by using BrainLab Smartbrush software (Brainlab, Munich, Germany). Pre-operative MRI scans were obtained within 24 h prior to resection, and post-operative scans were all obtained within 72 h post-resection. Total contrast-enhancing (CE) tumor volumes were measured at both pre-operative and post-operative time points. The total CE tumor volume was measured on T1-weighted post-contrast images, and the non-enhancing tumor volume was measured on T2 or FLAIR sequences. Manual segmentation was performed with region-of-interest analysis “painting” inclusion regions based on fluid-attenuated inversion-recovery (FLAIR) sequences from pre- and post-operative MRI scans to quantify tumor volume. Extent of Resection (EOR) was calculated as: (pre-operative tumor volume - post-operative tumor volume)/pre-operative tumor volume x 100%. Manual segmentations were performed with tumor volumetric verified for accuracy after an initial training period. Volumetric measurements were made blinded to patients’ clinical outcomes. All patients in the cohort had available preoperative and postoperative MRI scans for analysis. To ensure that post-operative FLAIR signal was not surgically induced edema or ischemia, FLAIR pre- and post-operative MRIs were carefully compared alongside DWI sequences prior to including each region in the volume segmentation^35^. HFC voxels with T1 post gadolinium contrast enhancing tumor were considered HFC-positive for survival analysis.

### Language Assessments

One to two days prior to tumor resection, patients underwent baseline language evaluation which consisted of naming pictorial representations of common objects and animals (Picture Naming, PN) and naming common objects and animals via auditory descriptions (Auditory Naming, AN). Visual picture naming (PN) and auditory stimulus naming (AN) testing were used given their known significance and clinical correlation with outcomes in clinical patient population ^57, 58^. The correct answers for these tasks (delivered on a laptop with a 15-inch monitor [60 Hz refresh rate] positioned two feet away from the seated patient in a quiet clinical setting) were matched on word frequency (i.e. commonality within the English language) using SUBTLEX_WF_ scores provided by the Elixcon project and content category. Task stimuli were randomized and presented using PsychToolbox. Task order was randomly selected by the psychometrist for each subject. Slides were manually advanced by the psychometrist either immediately after the subject provided a response or after six seconds if no response was given. The tasks were scored on a scale from 0 to 4 by a trained clinical research coordinator who was initially blinded to all clinical data (including imaging studies). No participants had uncorrectable visual or hearing loss. Details of the administration and scoring of auditory and picture naming language tasks can be found in prior studies^21, 39, 59^.

### Statistical analyses

Statistical tests were conducted using Prism (GraphPad) software unless otherwise indicated. Unpaired, two-tailed Student’s *t*-test was used to detect significance. A level of *P*<0.05 was used to designate significant differences. Two-tailed log rank analyses were used to analyze statistical significance of Kaplan–Meier survival curves. Statistical analyses for proteomic and RNA-seq data are described above in the respective sections.

### Data Availability

All data in the article are available from the corresponding author upon request.

## Acknowledgements

This study was supported by the NIH grants K08NS110919 and P50CA097257, the Robert Wood Johnson Foundation grant 74259, the UCSF LoGlio Collective, and the Sheri Sobrato Brisson Brain Cancer Fund to S.H-J.; Sullivan Brain Cancer Fund to S.K.; NIH grants F30CA246808 and T32GM007618 to A.C; the NIH grant K99CA252001 to H.S.V.; the NIH grant R01NS100440 to S.N.; the NIH grant R00DC013828 to D.B.; NIH grants R01NS092597 and DP1NS111132 and a Robert J. Kleberg, Jr. and Helen C. Kleberg Foundation grant to M.M.; American Brain Tumor Association grant MSSF1900021 and Lucien Rubinstein Award to N.A.; and the UCSF Physician Scientist Scholar Program, the UCSF Wolfe Meningioma Program Project, NIH grant K08CA212279 to D.R. The authors thank Joanna Phillips, Anny Shai and the staff of the UCSF Brain Tumor Center Biorepository and Pathology Core. We thank Applied Biomics Inc. for 2D-DIGE Proteomics analysis. The authors would like to acknowledge the staff within the Biological Imaging Development CoLab (BIDC) at UCSF for confocal microscope and Imaris software support. We thank Dr. Tomoko Ozawa and Ms. Raquel Santos of the UCSF Brain Tumor Center Preclinical Therapeutics Core. We thank Kenneth Probst for illustrations.

## Author contributions

S.K. (Krishna) and S.H-J. designed, conducted, and analyzed experiments. M.M. contributed to study design, data analysis, and manuscript editing. A.C. and D.R. contributed to bulk and single cell transcriptomic analyses. K.S. contributed to in vitro organoid experiments. L.N. contributed to electron microscopy data acquisition and analyses. S.K. (Kakaizada) and N.A. contributed to language and cognitive assessments. A.L. and A.A. contributed to human survival analyses. C.C. and R.S. contributed to quantitative imaging analysis. A.E., A.F. and S.N. contributed to MEG data acquisition and quantification. H.S.V. contributed to orthotopic xenografting. D.B. contributed to intraoperative electrocorticography analysis. S.K. (Krishna) and S.H-J. wrote the manuscript. S.H-J. conceived of the project and supervised all aspects of the work.

## Materials and Correspondence

All request for materials and correspondence may be sent to

---

**Extended Data Fig. 1.**
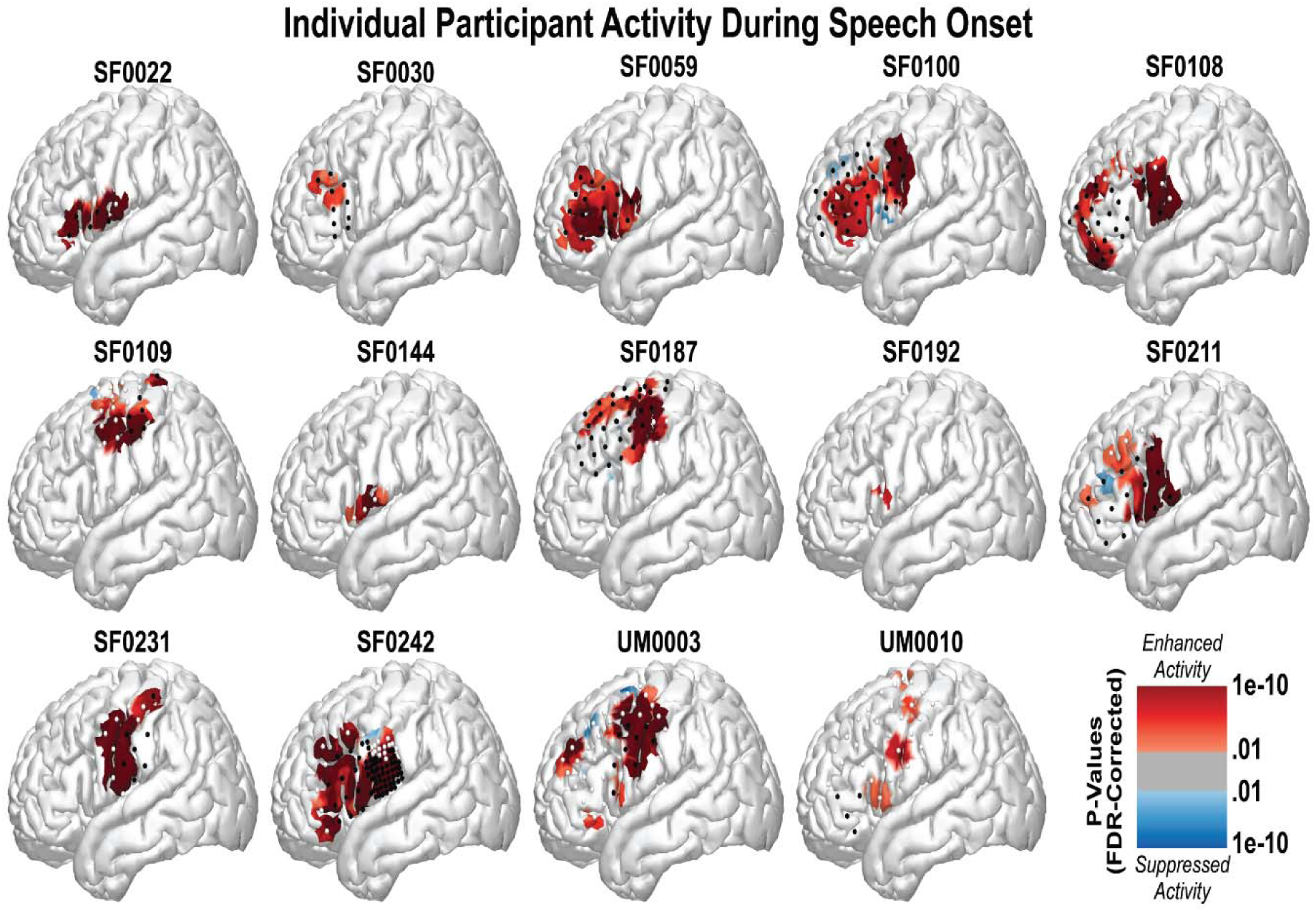
Individual cognitive task-related neuronal activity from a cohort of 14 adult patients with cortically projecting glioma infiltration in the lateral prefrontal cortex.

**Extended Fig. 2.**
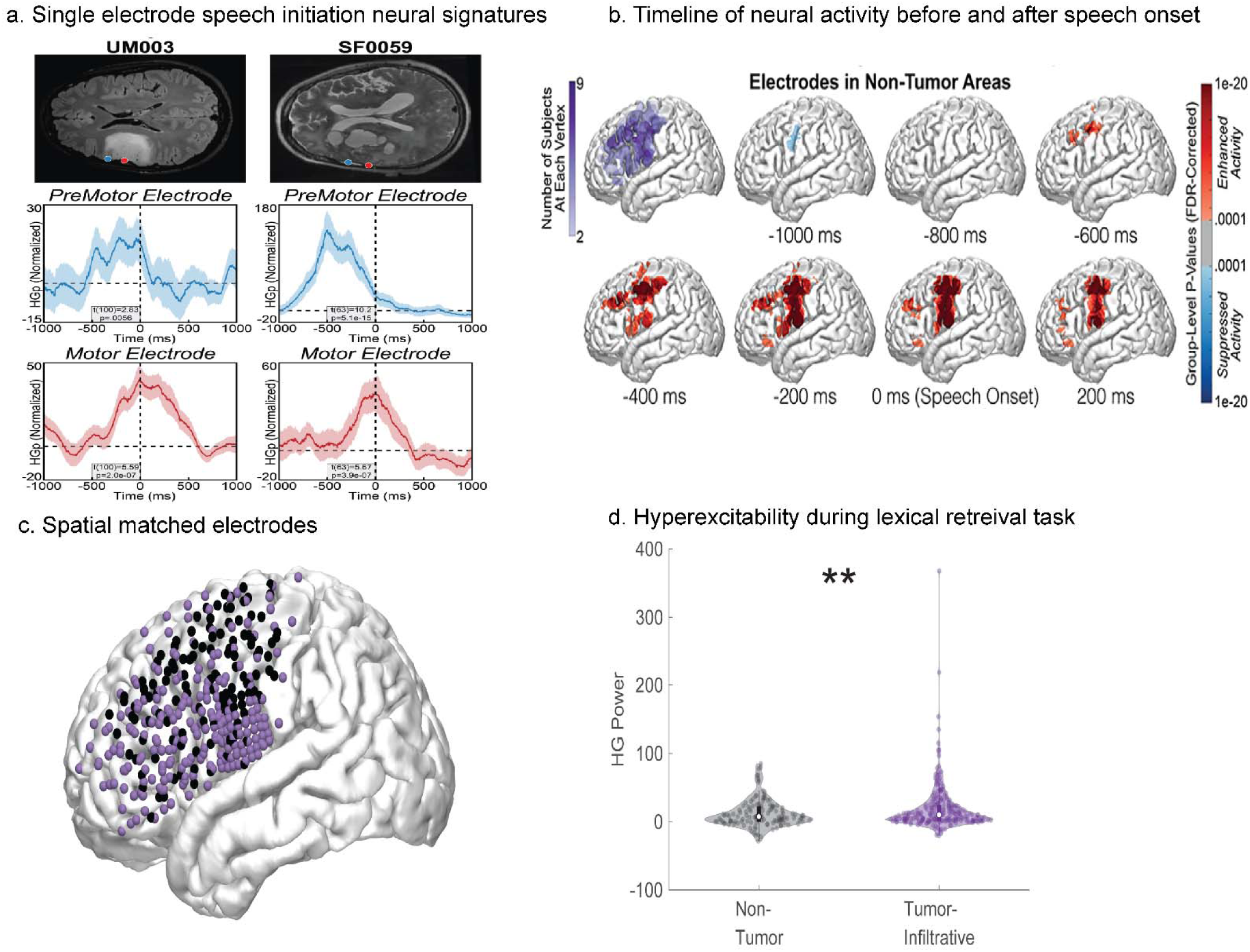
a, High gamma power (HGp) recording from single electrodes overlying tumor-infiltrated regions of brain. b, Reconstructed time series of high gamma neural activity from non-tumor electrodes demonstrating expected neural activity within lateral prefrontal cortex. c, d, Pair-match of each cortical electrode array within tumor-infiltrated cortical electrodes prior to speech onset.

**Extended Fig. 3.**
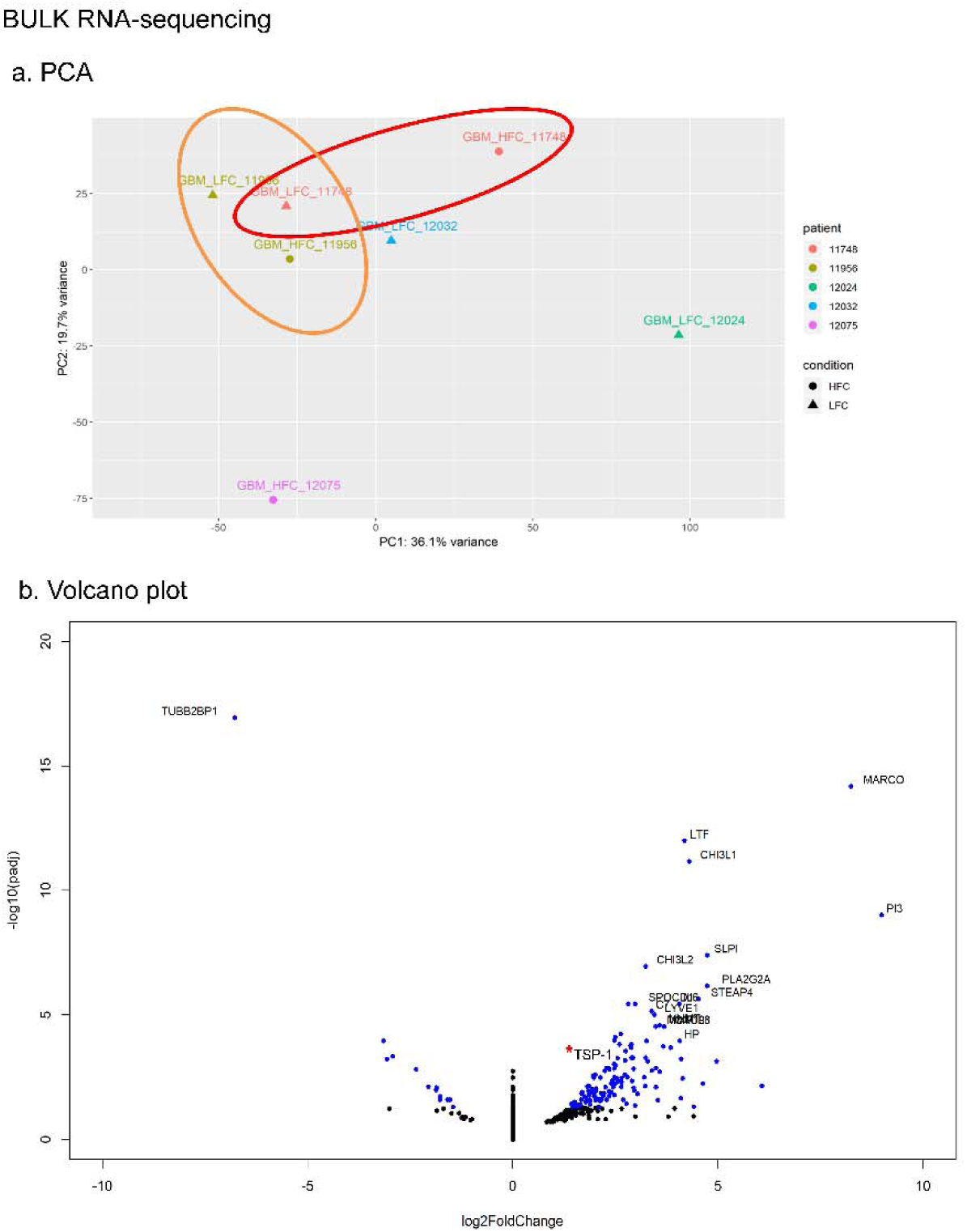
Bulk RNA-sequencing from primary patient-derived HFC and LFC tumor samples. a, Unsupervised principal component (PCA) analysis of bulk RNA sequencing data obtained from IDH-WT glioblastoma patient primary biopsy HFC (n = 3) and LFC (n = 4) tumor samples. b, Volcano plots of IDH-WT glioblastoma samples revealed 144 differentially expressed genes between HFC and LFC tumor regions. The blue dots represent all differentially expressed genes, where differential expression is defined by the parameters: adjusted p-value < 0.05 and absolute log2fold change > 1.

**Extended Fig. 4.**
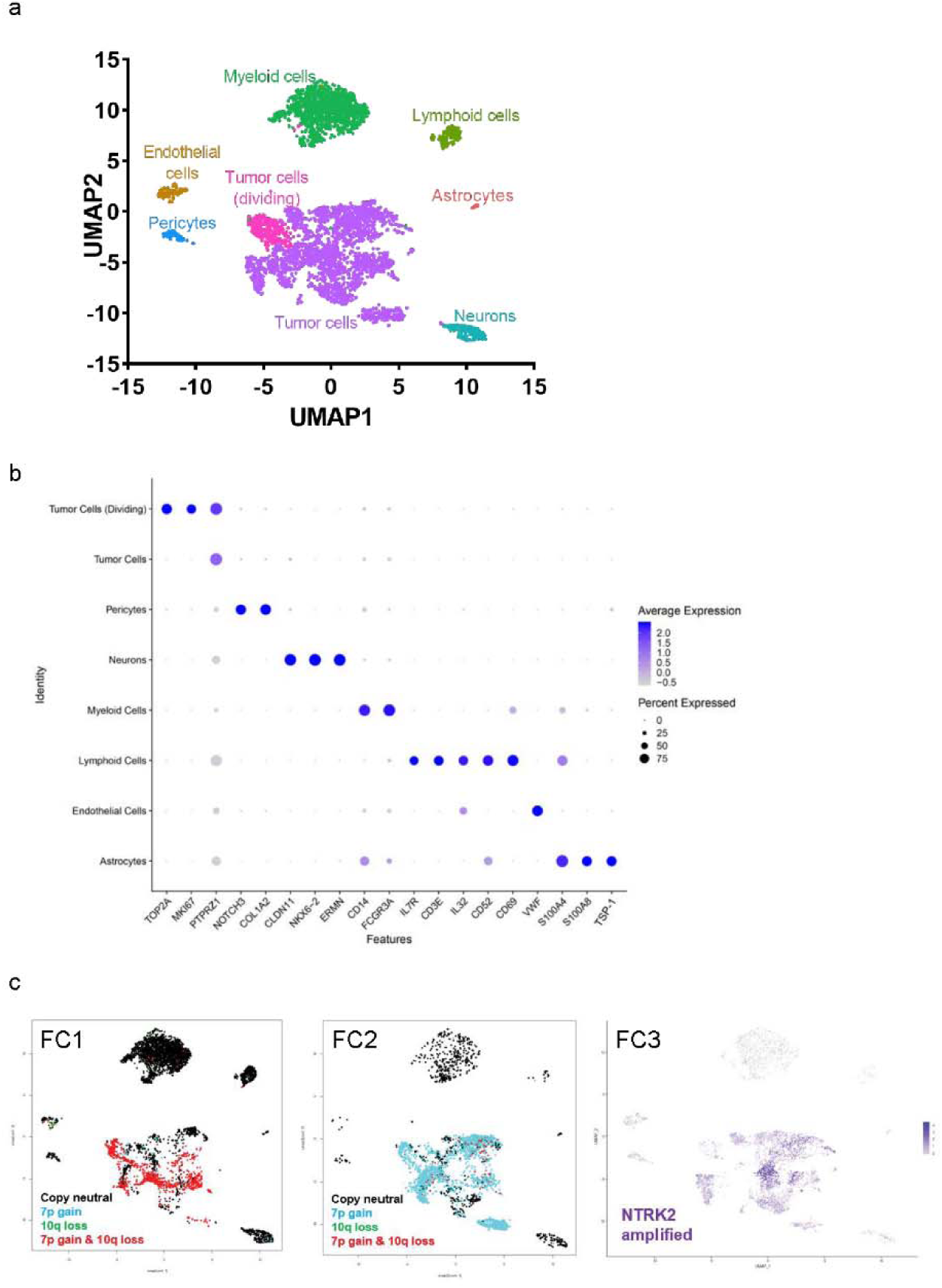
Single-cell RNA-sequencing from primary patient-derived HFC and LFC tumor samples. a, Single cell RNA transcriptomic profile UMAP confirms distinct cell populations including non-tumor astrocytes and neurons. b, Gene enrichment profile used to identify each of the UMAP cell populations. c, Tumor cell validation using copy number variant assessment on three matched pairs of HFC and LFC samples from FC1 (SF0200), FC2 (SF0209), and FC3 (SF0213) IDH-WT glioblastoma patients. Trisomy 7 and monosomy 10 co-occur in most cells in FC1.Trisomy 7 is an early event, while monosomy 10 is a late event in FC2. FC3 patient sample contains no copy number variation but has high level amplification of *NTRK2* gene.

**Extended Data Fig. 5.**
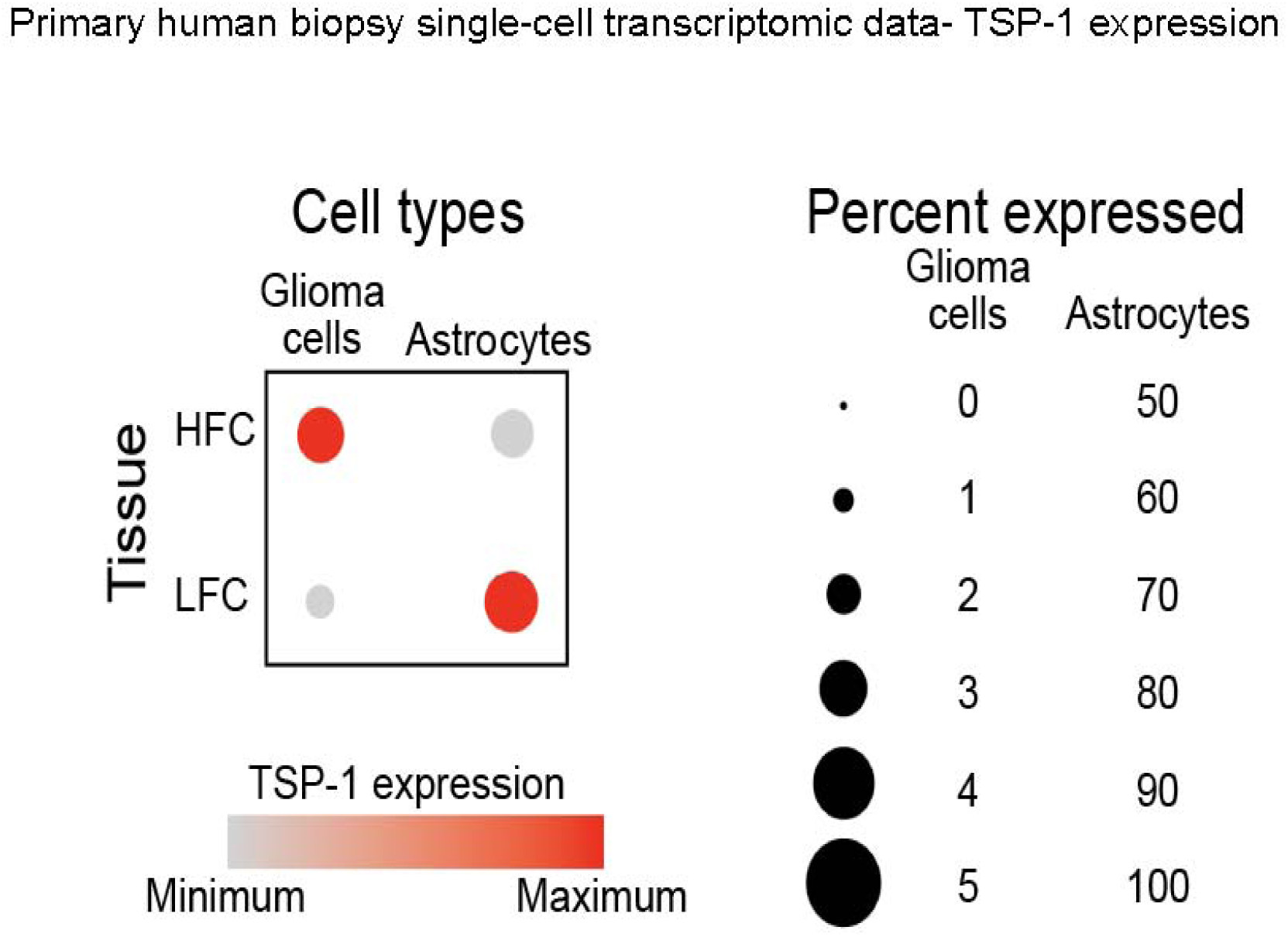
Primary human biopsy single-cell transcriptomic data showing increased thrombospondin-1 (*TSP-1*) expression in HFC high-grade glioma cell population. Dot plots showing *TSP-1* expression and percentage (number of cells expressing the gene) of *TSP-1*- positive cells in tumor cells and non-tumor astrocyte populations in HFC and LFC samples (n = 3 per group). Out of the total HFC tumor cells (n = 5325, 3 patients), 157 cells are *TSP-1* positive accounting for a percentage of 2.95, while only 1.59% (51 cells out of a total of 3212 LFC tumor cells [n = 3 patients]) express *TSP-1*. However, in the non-tumor astrocyte population, the number of *TSP- 1*-positive cells are higher in LFC (n = 34 out of a total of 41 astrocytes, accounting for 82.9%) compared to HFC (n = 15 out of a total of 20 astrocytes, accounting for 75%) samples.

**Extended Data Fig. 6.**
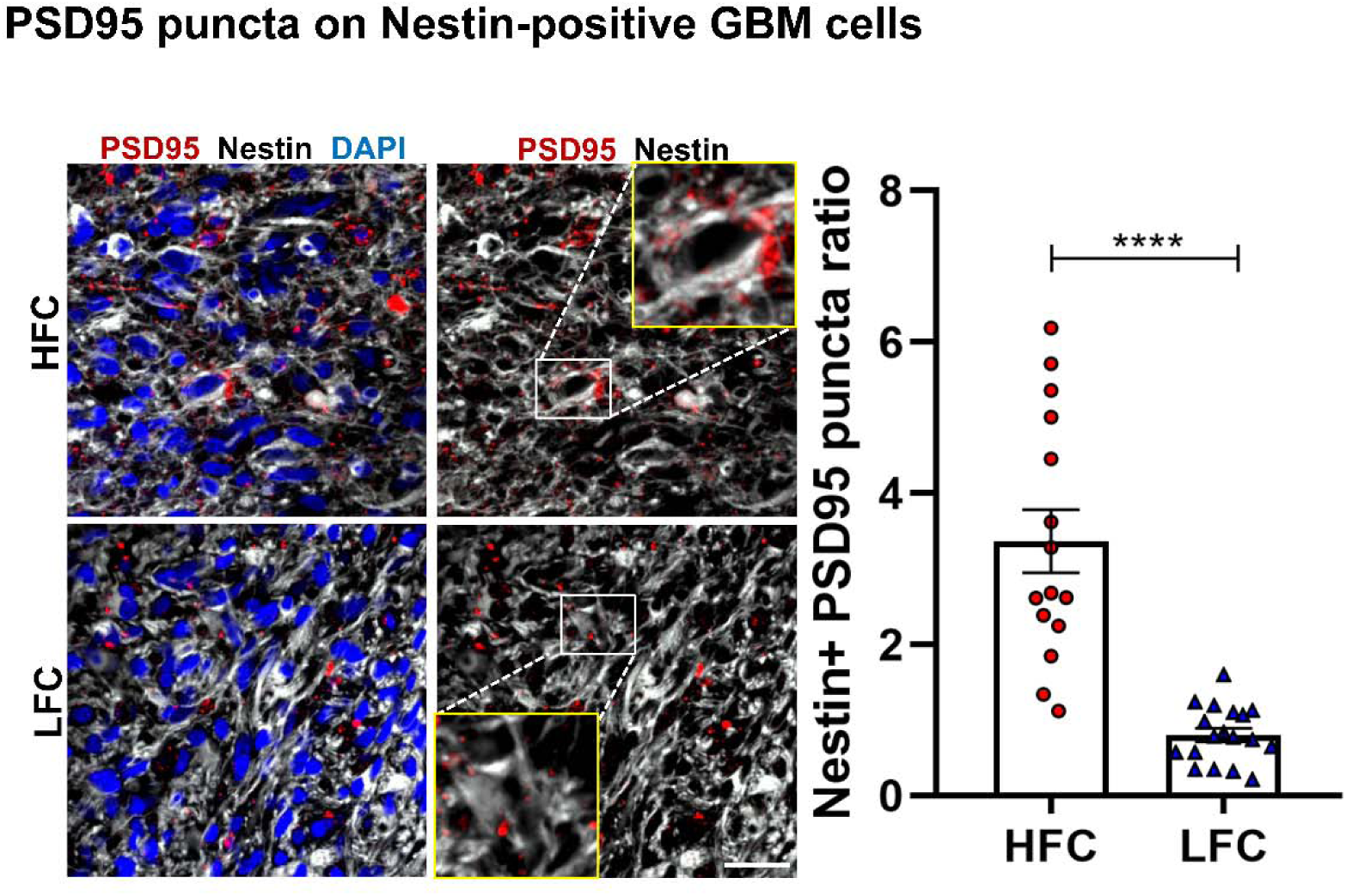
Tumor cells in HFC regions has increased postsynaptic marker (PSD95) expression compared to LFC. Ratio of nestin-positive postsynaptic PSD95 puncta count (calculated by dividing the total number of PSD95 puncta on nestin-positive tumor cells to the total number of cells stained with DAPI in 135 x 135-μm field areas) in GBM-HFC/ LFC tissue samples (HFC vs. LFC: 3.37 ± 0.42 vs. 0.81 ± 0.09); n = 15-18 sections per group, n =3 per group). Red, PSD95 (postsynaptic puncta); white, nestin (tumor cells); blue, DAPI. Scale bar, 10 µm. Inset shows zoomed-in view of PSD95 puncta on HFC/LFC tumor cells (positive for nestin). Scale bar, 3 µm. Data presented as mean ± s.e.m. P value determined by two-tailed Student’s t-test. \*\*\*\**P* < 0.0001.

**Extended Data Fig. 7.**
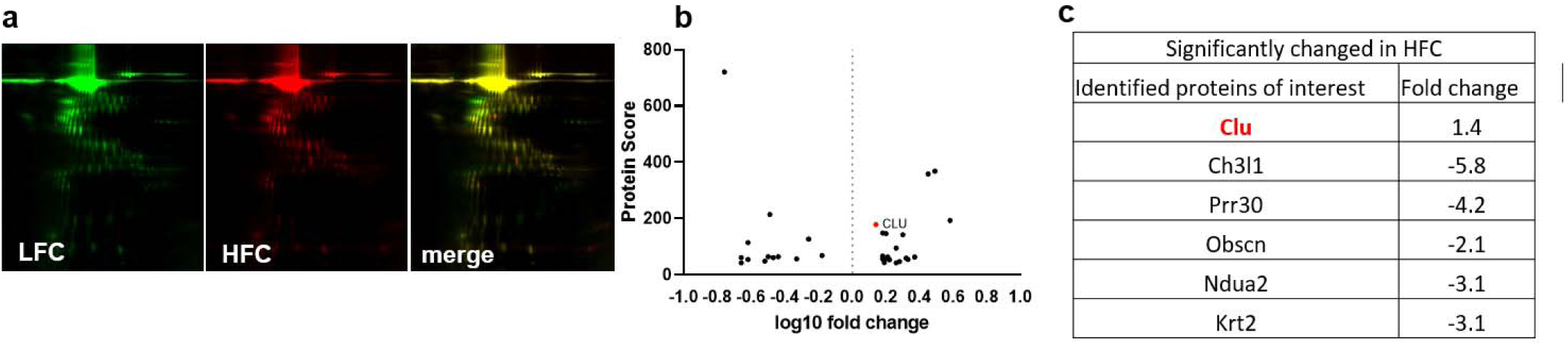
Proteomics analysis of secreted factors in HFC/LFC glioma cell culture. a, Proteomics analysis showing representative two-dimensional gel electrophoresis separating proteins in LFC (green) and HFC (red) by size (vertical axis) and charge (horizontal axis); merged image (right-most panel) with the red and green spots representing the up- and down-regulated proteins, respectively. Clusterin (CLU) was identified as a protein of interest with significant upregulation in HFC compared to LFC samples. b, Volcano plot showing protein score data of differentially regulated proteins in HFC compared to LFC samples. CLU is highlighted. c, List of candidate proteins of interest identified from proteomic analyses based on their established role in neurogenesis or tumorigenesis.

**Extended Data Fig. 8.**
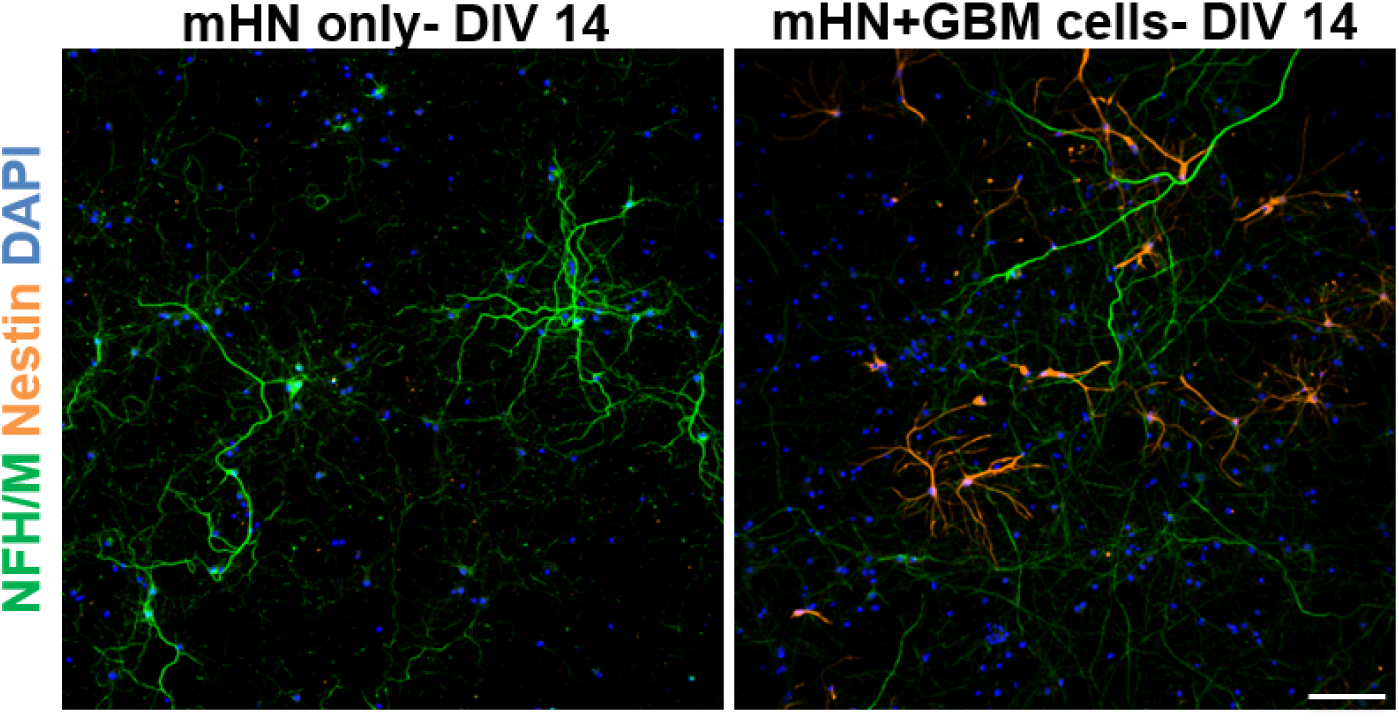
Nestin specifically labels tumor cells in glioma-neuron co-culture. Neurofilament (heavy and medium chains) and nestin antibodies used as specific markers to label mouse hippocampal neurons and HFC/LFC tumor cells, respectively in glioma-neuron co-culture. Left panel: mouse hippocampal neurons alone in culture for 14 days only express neurofilament (green) and not nestin (orange). Right panel: Nestin (orange) expression in GBM cells co-cultured with neurofilament (green) labelled mouse hippocampal neurons for 14 days. Scale bar, 100 µm.

**Extended Data Fig. 9.**
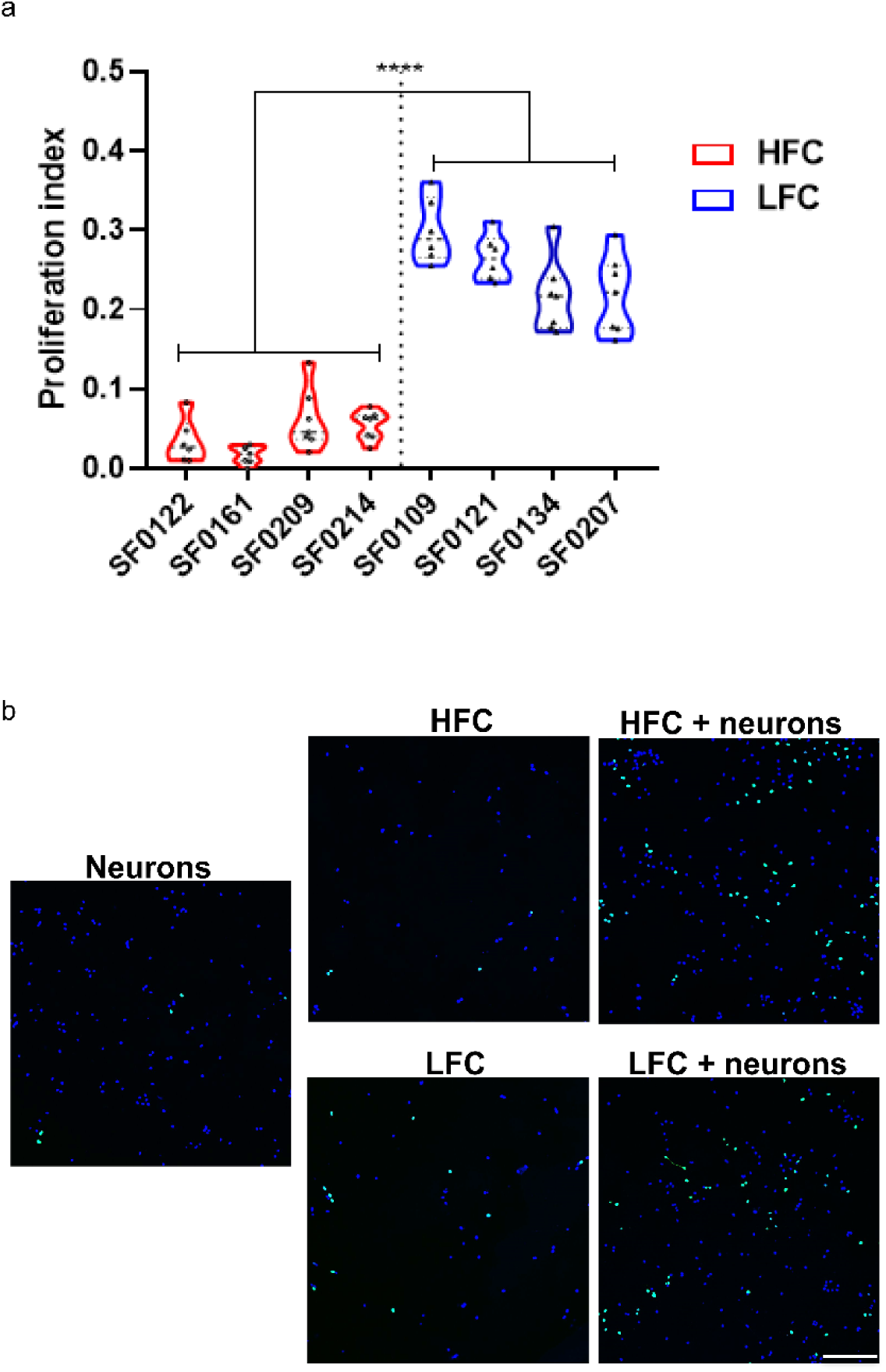
Glioma cells from HFC regions acquire proliferative phenotype in the presence of neurons. Primary patient glioma cells from HFC regions illustrate marked increase in proliferation when co-cultured with mouse hippocampal neurons. a, Quantification of proliferation indices of HFC (n = 4) and LFC (n = 4) glioma cells from individual patient lines, determined by quantifying the fraction of EdU labeled cells/DAPI labeled cells. b, Representative confocal images illustrating proliferating HFC and LFC glioma cells (EdU^+^, green) in the absence or presence of mouse hippocampal neurons (72h co-culture). Scale bar, 100 µm. Data presented as mean ± s.e.m. *P* value determined by two-tailed Student’s t-test. \*\*\*\**P* < 0.0001.

**Extended Data Fig. 10.**
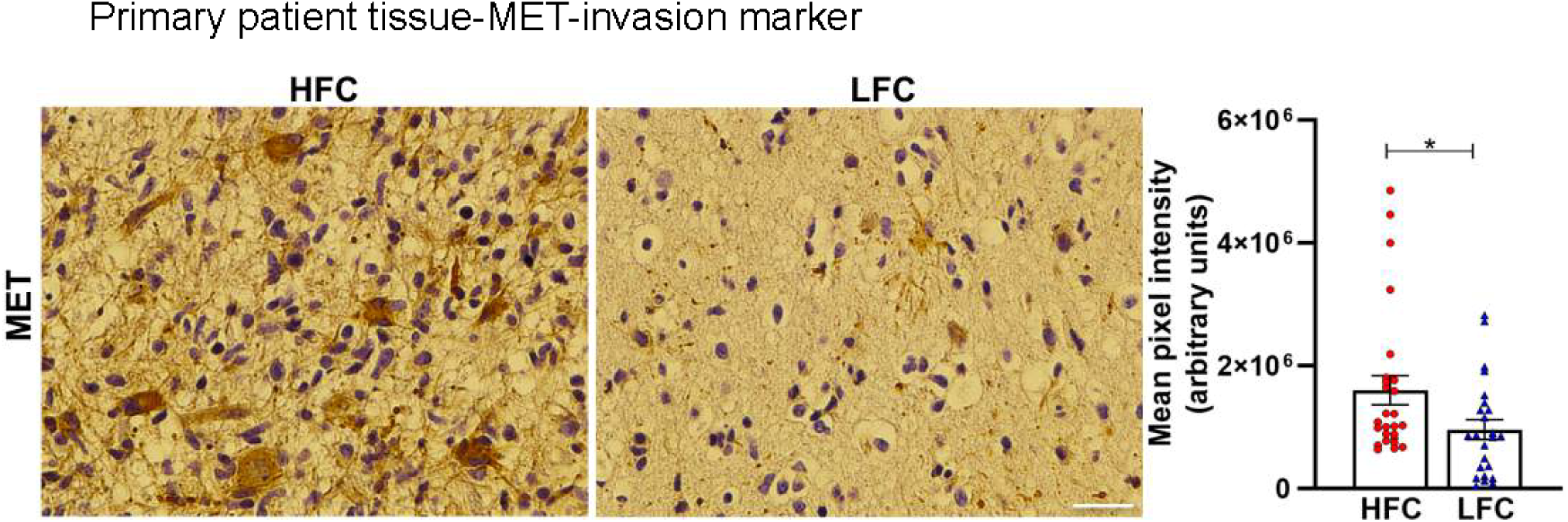
Site-directed biopsy tissues from HFC regions show increased expression of the invasive marker (MET) compared to LFC samples. Representative immunohistochemistry images of MET staining in HFC and LFC tissues demonstrate increased tissue level protein expression (HFC vs. LFC: 1,604,313.31 ± 235,778.38 vs. 965,734.39 ± 161,979.86, n = 4-5 images/sample, n = 4-5/group). Data presented as mean ± s.e.m. *P* values determined by two-tailed Student’s t-test. \**P* < 0.05.

**Extended Data Fig. 11.**
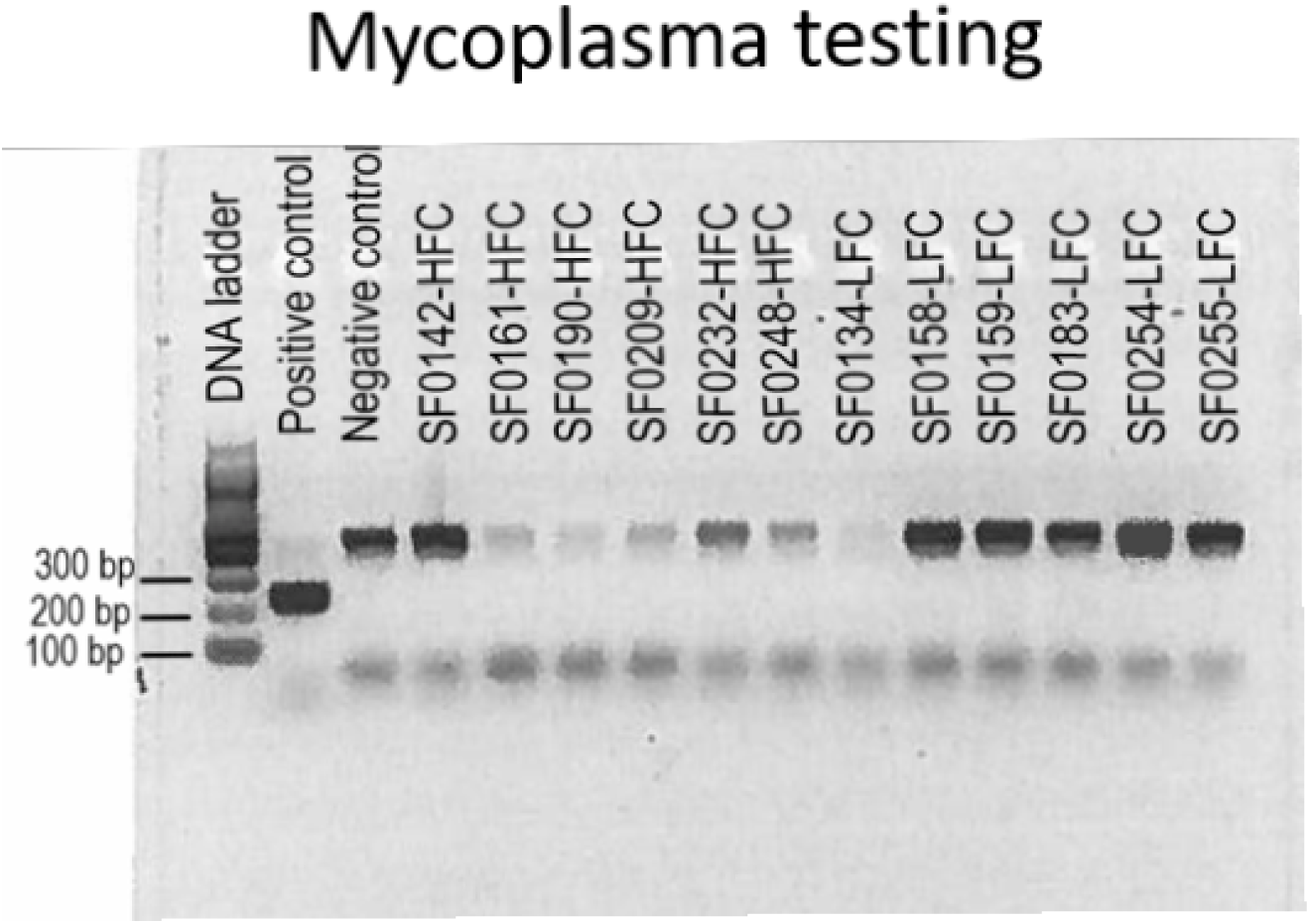
Mycoplasma screening of primary patient-derived cell lines. Cell cultures tested for mycoplasma using a commercially available kit (PCR Mycoplasma Test Kit I/C, PromoCell, Heidelberg, Germany) shows absence of a positive band at ∼270bp. Tested primary patient-derived lines shows internal control DNA at ∼479 bp indicated a successfully performed PCR.

**Extended Data Table 1.** List of patient samples used for each experiment. *✓Patient-matched samples.

| Patient # | Human ECoG | RNA Transcriptomics | Human Tissue IHC+ IF | 1° Patient Cell Culture | Protein Analysis (Proteomics+ ELISA) | PDX | Language Tasks | Human Survival |
| --- | --- | --- | --- | --- | --- | --- | --- | --- |
| 1 |  |  |  |  |  |  |  | ✓ |
| 2 |  |  |  |  |  |  |  | ✓ |
| 3 |  |  |  |  | ✓ |  | ✓ | ✓ |
| 4 |  |  |  |  |  |  | ✓ |  |
| 5 |  |  |  |  |  |  | ✓ | ✓ |
| 6 |  |  |  |  |  |  | ✓ | ✓ |
| 7 | ✓ |  |  |  |  |  |  |  |
| 8 |  |  |  |  | ✓ |  | ✓ | ✓ |
| 9 |  |  |  |  |  |  |  | ✓ |
| 10 | ✓ |  |  |  |  |  |  |  |
| 11 |  |  |  |  | ✓ |  | ✓ | ✓ |
| 12 |  |  |  |  |  |  |  | ✓ |
| 13 |  |  |  |  |  |  |  | ✓ |
| 14 |  |  |  |  |  |  |  |  |
| 15 | ✓ |  |  |  |  |  |  |  |
| 16 |  |  |  |  |  |  |  | ✓ |
| 17 |  | *✓ | *✓ |  | ✓ |  |  |  |
| 18 |  |  |  |  | ✓ |  |  | ✓ |
| 19 |  |  |  |  |  |  | ✓ |  |
| 20 |  |  |  |  | ✓ |  |  |  |
| 21 |  |  |  |  | ✓ |  | ✓ | ✓ |
| 22 |  |  |  |  | ✓ |  | ✓ | ✓ |
| 23 |  |  |  |  |  |  |  | ✓ |
| 24 |  |  |  |  |  |  |  | ✓ |
| 25 | ✓ |  |  |  |  |  |  |  |
| 26 |  |  |  |  | ✓ |  |  | ✓ |
| 27 | ✓ |  |  |  |  |  |  |  |
| 28 | ✓ | *✓ | *✓ | *✓ | *✓ |  | ✓ | ✓ |
| 29 |  |  |  |  |  |  | ✓ | ✓ |
| 30 |  |  |  | *✓ | *✓ |  |  | ✓ |
| 31 |  | ✓ |  | ✓ |  |  |  | ✓ |
| 32 |  |  | ✓ | ✓ | ✓ | ✓ |  | ✓ |
| 33 |  | ✓ | ✓ | ✓ | ✓ |  | ✓ | ✓ |
| 34 |  | ✓ | ✓ | ✓ |  | ✓ | ✓ | ✓ |
| 35 |  |  |  | ✓ |  |  |  | ✓ |
| 36 |  |  |  |  | ✓ |  | ✓ | ✓ |
| 37 |  |  |  |  |  | ✓ |  | ✓ |
| 38 |  |  | ✓ |  |  |  |  | ✓ |
| 39 | ✓ |  |  |  |  |  |  |  |
| 40 |  |  |  |  |  |  | ✓ |  |
| 41 |  |  |  |  |  |  |  | ✓ |
| 42 |  |  |  |  | ✓ |  | ✓ | ✓ |
| 43 |  |  |  | ✓ |  |  |  | ✓ |
| 44 |  |  | ✓ | ✓ | ✓ |  |  |  |
| 45 |  |  | *✓ |  |  |  |  |  |
| 46 |  |  |  |  | ✓ | ✓ |  | ✓ |
| 47 |  |  |  |  |  |  |  | ✓ |
| 48 |  |  | ✓ | *✓ |  |  |  |  |
| 49 | ✓ |  |  |  |  |  |  |  |
| 50 |  |  |  | *✓ |  |  |  |  |
| 51 | ✓ |  |  |  |  |  |  |  |
| 52 |  |  |  | *✓ |  |  |  |  |
| 53 |  | *✓ | *✓ | ✓ |  |  |  |  |
| 54 |  |  |  |  | ✓ |  |  |  |
| 55 |  |  |  | ✓ |  |  |  |  |
| 56 |  | *✓ | *✓ | ✓ |  |  |  |  |
| 57 | ✓ |  |  |  |  |  |  |  |
| 58 |  | *✓ |  | ✓ |  |  |  |  |
| 59 |  |  | *✓ | ✓ |  |  |  |  |
| 60 | ✓ |  |  |  |  |  |  |  |
| 61 |  |  |  | ✓ |  |  |  |  |
| 62 |  |  |  | ✓ |  |  |  |  |
| 63 | ✓ |  |  |  |  |  |  |  |
| 64 |  |  |  |  |  | ✓ |  |  |
| 65 |  |  |  |  |  | *✓ |  |  |
| 66 |  |  |  |  |  | ✓ |  |  |
| 67 |  |  |  |  |  |  |  | ✓ |
| 68 | ✓ |  |  |  |  |  |  |  |
| 69 | ✓ |  |  |  |  |  |  |  |
| 70 |  |  |  |  |  |  |  | ✓ |
| 71 |  |  |  |  |  |  |  | ✓ |
| 72 |  |  |  |  |  |  |  | ✓ |
| 73 |  |  |  |  |  |  |  | ✓ |
| 74 |  |  |  |  |  |  |  | ✓ |
| 75 |  |  |  |  |  |  |  | ✓ |
| 76 |  |  |  |  |  |  |  | ✓ |
| 77 |  |  |  |  |  |  |  | ✓ |
| 78 |  |  |  |  |  |  |  | ✓ |
| 79 |  |  |  |  |  |  |  | ✓ |
| 80 |  |  |  |  |  |  |  | ✓ |
| 81 |  |  |  |  |  |  |  | ✓ |
| 82 |  |  |  |  |  |  |  | ✓ |
| 83 |  |  |  |  |  |  |  | ✓ |
| 84 |  |  |  |  |  |  |  | ✓ |
| 85 |  |  |  |  |  |  |  | ✓ |
| 86 |  |  |  |  |  |  |  | ✓ |
| 87 |  |  |  |  |  |  |  | ✓ |
| 88 |  |  |  |  |  |  |  | ✓ |
| 89 |  |  |  |  |  |  |  | ✓ |
| 90 |  |  |  |  |  |  |  | ✓ |
| 91 |  |  |  |  |  |  |  | ✓ |
| 92 |  |  |  |  |  |  |  | ✓ |
| 93 |  |  |  |  |  |  |  | ✓ |
| 94 |  |  |  |  |  |  |  | ✓ |
| 95 |  |  |  |  |  |  |  | ✓ |
| 96 |  |  |  |  |  |  |  | ✓ |
| 97 |  |  |  |  |  |  |  | ✓ |
| 98 |  |  |  |  |  |  |  | ✓ |
| 99 |  |  |  |  |  |  |  | ✓ |
| 100 |  |  |  |  |  |  |  | ✓ |
| 101 |  |  |  |  |  |  |  | ✓ |
| 102 |  |  |  |  |  |  |  | ✓ |
| **Total**<br>**N** | **14** | **13** | **18** | **24** | **20** | **8** | **16** | **66** |

**Extended Table 2.** Bulk RNA-seq based detection of the significant differentially expressed genes in high and low connectivity regions with log fold changes greater than 1 and an adjusted *P*-value less than 0.05.

| Gene Name | Log2 Fold-change | *P*-value | Adjusted *P*-value |
| --- | --- | --- | --- |
| TSP-1 | 2.89 | 1.02E-06 | 5.26E-04 |
| TGFBI | 2.02 | 5.08E-05 | 9.84E-03 |
| NTNG1 | 1.59 | 3.30E-04 | 3.91E-02 |
| SYNPO | 1.46 | 4.42E-04 | 4.91E-02 |

A negative fold change means that the gene is overexpressed in low connectivity tumor regions.

**Extended Data Table 3.** Single cell sequencing of three matched pairs of HFC and LFC samples from IDH-WT glioblastoma patients.

| Sample | Cells Before | Cells After |
| --- | --- | --- |
| HFC1 (SF0200) | 1277 | 967 |
| LFC1 (SF0200) | 6671 | 4277 |
| HFC2 (SF0209) | 2999 | 2529 |
| LFC2 (SF0209) | 666 | 549 |
| HFC3 (SF0213) | 4619 | 3170 |
| LFC3 (SF0213) | 3914 | 2239 |
| **Total** |  | **13,731** |

Cells before and after QC/filtering. Filters chosen: $500 < \text{nFeature\_RNA} < 10,000$ & $\text{percent.mt} < 0.2$ .

**Extended Data Table 4.** Demographics and clinical summary of patient cohorts used for survival data.

|  | **HFC-** | **HFC+** | ***P*-value** |
| --- | --- | --- | --- |
| No. of patients | 41 | 25 | - |
| Mean age (+/- SEM) | 60.6 (1.74) | 60.1 (2.52) | 0.82 |
| Gender (% male) | 53.7 | 60.0 | 0.61 |
| Tumor volume, cm<sup>3</sup> (+/- SEM) | 21.4 (4.47) | 30.3 (4.52) | 0.19 |
| Extent of resection (% , +/- SEM) | 97.9 (50) | 96.8 (86) | 0.25 |
| IDH (% WT) | 41 (100) | 25 (100) | 1 |
| MGMT (% methylation) | 58.5 | 72.0 | 0.27 |

## Supplementary Video 1

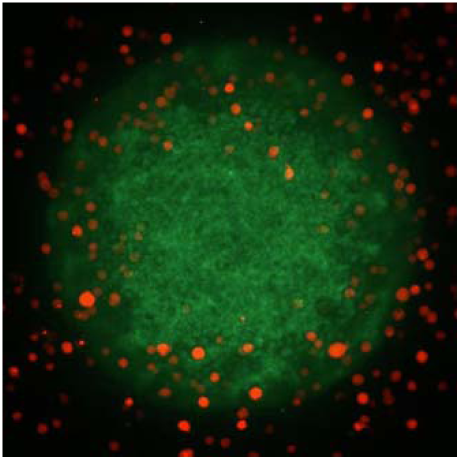

**3D co-culture of HFC glioma cells with induced neuron organoids**. Live cell imaging of HFC glioma cells (labelled with RFP) in co-culture for 12h with induced neuron organoids (labelled with GFP).

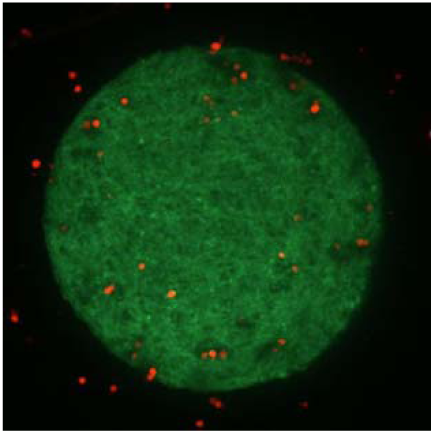

**3D co-culture of LFC glioma cells with induced neuron organoids**. Live cell imaging of LFC glioma cells (labelled with RFP) in co-culture for 12 h with induced neuron organoids (labelled with GFP).

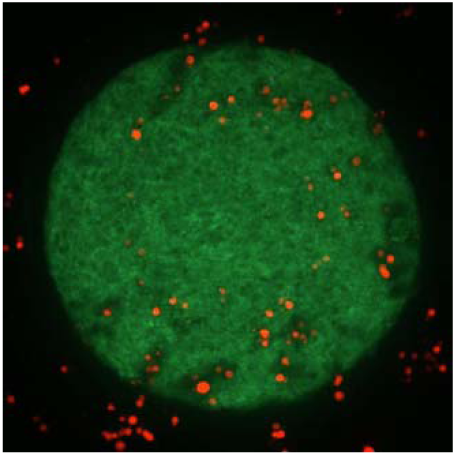

**3D co-culture of LFC glioma cells with induced neuron organoids in the presence of TSP-1.** Live cell imaging of HFC glioma cells (labelled with RFP) in co-culture for 12 h with induced neuron organoids (labelled with GFP). TSP-1 was exogenously applied at a dose of 5 µg/ml to the culture media.

